# Maternal Western-Style Diet Impairs Bone Marrow Development and Drives a Hyperinflammatory Phenotype in Hematopoietic Stem and Progenitor Cells in Fetal Rhesus Macaques

**DOI:** 10.1101/2021.04.26.441556

**Authors:** Suhas Sureshchandra, Chi N. Chan, Jacob J. Robino, Lindsay K. Parmelee, Michael J. Nash, Stephanie R. Wesolowski, Eric M. Pietras, Jacob E. Friedman, Diana Takahashi, Weining Shen, Jon D. Hennebold, Devorah Goldman, William Packwood, Jonathan R Lindner, Charles T. Roberts, Benjamin J. Burwitz, Ilhem Messaoudi, Oleg Varlamov

## Abstract

**Background:** Maternal obesity adversely impacts the in utero metabolic environment and offspring’s health, but its effect on fetal hematopoiesis and immune cell development remains incompletely understood, particularly in models that resemble human development.

**Methods:** We studied gestational day 130-135 fetuses derived from rhesus macaque dams chronically exposed to a high-fat Western-style diet (WSD) or a low-fat control diet. Fetal immune cell phenotypes and fetal bone marrow architecture and hematopoietic stem and progenitor cell (FBM HSPC) function were examined using bone computed tomography, histology, flow cytometry, single-cell RNA-sequencing, and HSPC transplantation assays.

**Findings:** Maternal WSD induced premature FBM cavity opening and a codominant increase in the number of FBM adipocytes. Furthermore, a maternal WSD induced a proinflammatory transcriptional response in FBM HSPCs. FBM macrophages from the WSD group exhibited heightened proinflammatory responses to toll-like receptor agonist stimulation. Maternal WSD exposure suppressed the expression of genes required for B-cell development and decreased the frequencies of FBM B-cells. Finally, maternal WSD led to poor engraftment of FBM HSPCs in nonlethally irradiated immunodeficient NOD/SCID/IL2rγ^-/-^ mice.

**Interpretations:** Maternal WSD impairs FBM development, drives a hyperinflammatory phenotype, and induces functional and differentiation impairment in FBM HSPCs in a translationally relevant nonhuman primate model.

**Funding:** National Institute of Health

**RESEARCH IN CONTEXT:** *Evidence before this study:* Maternal obesity is associated with increased risk of infections and proinflammatory disease in offspring. The translationally-relevant rhesus macaque model was utilized to address the effects of maternal obesogenic diet on fetal hematopoietic and immune cell development.

*Added value of this study:* We assessed changes in fetal immune cell phenotypes and fetal hematopoietic stem and progenitor cell function using immunohistochemistry, flow cytometry, single-cell RNA sequencing, and transplantation assays. We determined that chronic consumption of a maternal obesogenic diet induced the development of adipogenic and proinflammatory environments in the fetal bone marrow. Additionally, we detected the impairment in B-cell differentiation program in fetal hematopoietic stem and progenitor cells.

*Implications of all the available evidence:* These data demonstrate that maternal obesogenic diet modulates fetal hematopoietic development and could impact the offspring’s immune system, including proinflammatory phenotype and a decline in B-cell function.

## INTRODUCTION

Pre-pregnancy (pregravid) obesity is associated with adverse outcomes for both mother and fetus, with long-term health consequences for the offspring ^1,2^. Additionally, maternal obesity is associated with increased risk of infections in offspring, including necrotizing enterocolitis, sepsis, and severe respiratory syncytial virus infection that necessitates admission to the neonatal intensive care unit ^3–5^. Studies using rodent and nonhuman primate (NHP) models demonstrated that pregravid maternal obesity leads to aberrant inflammatory responses in the offspring ^3,6–9^. While our understanding of the transgenerational impact of maternal obesity on the offspring’s immune system is beginning to emerge, the mechanisms that underlie the modulation of the fetal hematopoietic system and immune cell lineage specification during development are poorly understood. Furthermore, there is a paucity of studies that examined the impacts of maternal obesity on fetal hematopoiesis in models that resemble human development.

Hematopoiesis occurs in several waves that involve the migration of hematopoietic stem and progenitor cells (HSPCs) between different hematopoietic sites ^10^. Following the emergence of HSPCs in the aorta-gonad-mesonephros (AGM) during the first trimester, the fetal liver (FL) becomes the main hematopoietic niche for HSPC expansion ^11,12^. During the second and third trimesters, HSPCs migrate from the FL to the fetal bone marrow (FBM), representing an essential developmental stage associated with a transition to postnatal hematopoiesis ^10^. This emergence of HSPCs in the FBM coincides with the development of the FBM vascular system and bone ossification ^13,14^. Immune cell fate-mapping analyses and other techniques to assess cellular ontogeny have shed critical insight into the origins of tissue-resident macrophages ^15,16^. Early in gestation, tissue-resident macrophage progenitors are trapped in blood islands of the yolk sac until the circulatory system is established, at which point they colonize developing embryonic tissues, such as the FL and thymus, and maintain their proliferative capacity ^17,18^. As embryonic structures develop, the FBM becomes the major site where monocyte-derived macrophages, red blood cells, and B-lymphoid cells are produced throughout life ^15,19,20^. At this point, primitive yolk-sac derived macrophages are depleted in most tissues, aside from microglia, after being overrun by definitive HSPC-derived macrophages ^17,21^. Peripheral blood monocytes, when cued by environmental signals, can differentiate into macrophages ^22,23^. Both monocytes and tissue-resident macrophages are important for anti-microbial defense and tissue homeostasis ^15^. Therefore, alterations in the maternal environment may exert long-lasting effects on offspring immunity by reprogramming HSPC outputs in the FBM and the FL ^24^ via intrinsic (e.g., change in HSPC transcriptome) and extrinsic (e.g., change in a FBM niche) mechanisms; however, the molecular mechanisms remain poorly understood.

In the present study, we further explored these mechanisms by leveraging a rhesus macaque model, whose hematopoietic ^25–28^ and immune ^29^ systems are highly similar to that of humans. We analyzed immune system development in gestational day (GD) 130-135 fetuses derived from dams chronically exposed to a high-fat/calorie-dense Western-style diet (WSD) ^30^ or a low-fat/low-calorie control diet ^31,32^. Fetal immune cell phenotypes and fetal HSPC functional properties were examined using a combination of immunohistochemistry, flow cytometry, single-cell RNA-sequencing (scRNA-Seq), and HSPC transplantation assays (see experimental design in **Supplementary Figure S1A**). Here, we report for the first time that the chronic consumption of a maternal obesogenic WSD stimulates FBM adipogenesis and leads to a FBM proinflammatory state and functional impairment in differentiation of fetal HSPCs.

## METHODS

### Animal procedures

All animal procedures were approved by the Oregon National Primate Research Center (ONPRC) Institutional Animal Care and Use Committee and comply with the Animal Welfare Act and the APA Guidelines for Ethical Conduct in the Care and Use of Nonhuman Animals in Research. Dietary interventions, animal care, and fertility trial procedures have been previously reported ^31–34^. Before C-section, food was withheld for approximately 12 hours prior to the procedure. Water was not withheld. Animals were sedated with 8-20 mg/kg ketamine. Following sedation, a local block consisting of 0.8 ml bupivicaine (0.5%) combined with 0.2 ml lidocaine (1%) with epinephrine was placed intradermally at the incision site. Once the intravenous catheter was placed, the animal received 0.025-0.2 mg/kg hydromorphone intravenously. Animals were endotracheally intubated with an appropriately sized endotracheal tubes and general anesthesia was induced with 3% Isoflurane for 2-3 minutes or until the animal began to lose reflexes and heart rate slowed. Inhalant anesthesia was maintained at 1-2% isoflurane which was adjusted based on the physiologic parameters of the animal. Inhalant anesthetics were combined with 100% oxygen administered at a rate of 1-1.5 L/min. All surgical procedures were conducted by trained ONPRC Surgical Services Unit personnel under the supervision of surgical veterinarians in dedicated surgical facilities using aseptic techniques and comprehensive physiological monitoring. All animals had 22-gauge cephalic catheters placed and were intubated in dorsal recumbency with an appropriate size endotracheal tubes. Animals received intravenous fluids (generally Lactated Ringers Solution) at 10 ml/kg/hour; fluids were increased based on hydration status of animal or due to blood loss during surgery. The animal was positioned in dorsal recumbency, followed by sterile prep and draping. The abdomen was entered via 10-cm linear ventral midline laparotomy, followed by delivery and draping of the gravid uterus with moistened laparotomy sponges. The fetus was balloted to the fundic region, and a transverse hysterotomy was made and delivery of the fetus was completed. Fetuses were sedated with 10-20 mg/kg of ketamine and transported to the ONPRC Pathology Services Unit. Fetuses were anesthetized with sodium pentobarbital administered via the umbilical vessels at 25 mg/kg and euthanized by exsanguination. Fetal bone samples were collected and processed within 15 minutes of exsanguination.

### Bone immunohistochemistry and image analysis

Histological analyses were performed on paraffin-embedded sections. Specifically, fetal femurs and sterna were fixed in 4% PFA for 24 hours at 4^0^C, decalcified in 0.5 M EDTA for 4-7 days, and embedded in paraffin. Adjacent 5-μm paraffin sections were stained with hematoxylin and eosin to evaluate bone morphology and adipogenesis. 5-μm thick sternal sections were placed on glass slides and baked overnight at 60°C. Slides were deparaffinized and heat induced epitope retrieval was performed with the ACDBio Target Retrieval Reagent in a Biocare DeCloaking Chamber at 110°C for 30 minutes. Slides were then washed in dH_2_O and TBS-T before incubation overnight at room temperature with a rabbit anti-CD34 antibody (HPA036723, Millipore Sigma), a mouse anti-CD20 antibody (M0755, clone L26, Dako, Santa Clara CA, USA), or for macrophages 1 hour at room temperature with a mixture of mouse anti-CD68 (CM033B, clone KP1, Biocare, Pacheco, CA, USA) and mouse anti-CD163 antibody (catalogue number 11458, Invitrogen). Slides were then washed in TBS-T and treated with 3% H_2_O_2_ in PBS for 5 minutes. Detection of the primary antibodies was performed with Gold Bridge International Labs Polink 1 HRP Detection System against Rabbit IgG (D13-110), or Mouse IgG (D12-110) (CD34/CD20) or with the Polink 2 HRP Detection System against mouse IgG (D37-110) (CD68/163 combination). Staining was visualized with the chromogen 3,3′-Diaminobenzidine (DAB) and the sections were counterstained with hematoxylin and coverslipped. Whole slide scan images were captured using the Leica AT2 System at 40X. Quantitative tissue analysis was performed with HALO v3.2 (Indica Labs, Albuquerque, NM, USA) using the analysis module “Multiplex IHC v3.0.4” to detect total and DAB-positive cell numbers. To detect the proximity of HSPCs and macrophages to adipocytes and sinusoids, the locations of sinusoids and adipocytes were annotated based on a subjacent section stained with hematoxylin and eosin stains. Then an Infiltration analysis was performed to quantify the number of DAB-stained cells within set distances from the annotated sinusoids.

### Bone micro computed tomography and image analysis

The proximal and distal halves of the fetal femurs were placed in separate Eppendorf tubes containing 1 mL PBS. CT data acquisition was performed on an Inveon micro SPECT/CT instrument (Siemens, Inc). A 360-degree collection arc with 1 degree of separation between steps and single exposure of 3 sec at 40kV and 300 μA at each step was used. A 0.5-mm aluminum filter of the X-ray source was employed. The CT detector was placed in the medium-high position and data were binned in a 2X2 pattern resulting in a voxel size of approximately 29 microns. Image reconstruction was performed using Inveon Research Workplace software (Siemens, Inc.). Setting included the Feldkamp reconstruction algorithm, a noise reduction setting of Slight, and a Shepp-Logan filter. A Rat setting was used for bean hardening. CT image series were processed using the BoneJ plugin for ImageJ. Specifically, the stacks of images were converted into binary and processed in BoneJ using built-in functions.

### Cell isolation

For FBM cell isolation, both fetal rhesus macaque tibias were crushed manually with a ceramic mortar and pestle and the cell suspension was filtered through a 30-μm nylon cell strainer (Thermo Fisher Scientific, Waltham, MA, USA) in 20 ml ice-cold *X-Vivo* media (Lonza, Basel, Switzerland). FBM mononuclear cells were isolated by centrifugation for 30 minutes at room temperature on a Ficoll gradient (StemCell Technologies, Vancouver, BC, Canada) and red blood cells were removed using the RBC lysis buffer (StemCell Technologies). Mononuclear cells were washed three times with PBS and CD34+ cells were isolated as described ^35^ using rhesus macaque-specific antibodies to CD34 (clone 563; BD Pharmingen, San Jose, CA, USA) and the anti-PE MACS magnetic bead system (Miltenyi Biotec, Bergisch Gladbach, Germany) with the supplied reagents. CD34+ HSPC and CD34-flow-through cell fractions were cryopreserved in several aliquots using CryoStor CS10 (Stem Cells Technologies). FL mononuclear cells were obtained via perfusion and collagenase digestion of a portion of the liver from control and WSD fetuses, as reported previously ^36^. Briefly, hepatocytes were removed from the total mixture of digested cells by centrifugation at 100 x g for 5 minutes. The supernatant, containing all non-parenchymal cells, was filtered and spun at 800 x g for 10 minutes at 4° C to pellet non-parenchymal cells. Resulting cell pellets were resuspended in 24% Histodenz (Sigma-Aldrich, St. Louis, MO, USA) and gradients were prepared and spun at 1500 x g for 20 minutes at 4° C with the brake turned off. Cells at the resulting interface (liver mononuclear cells) were collected and washed, and aliquots were cryopreserved in 90% FBS/10% DMSO, stored in liquid N_2_, then thawed and used for flow cytometry analysis.

### Engraftment of HSPCs in NSG mice

The procedures were performed on 4-week-old female NSG mice (Jackson Laboratory, Bar Harbor, ME, USA; Strain: 005557 NOD.Cg-Prkdc Il2rg/SzJ) of at least 15 grams body weight. The NSG mice were maintained on regular rodent chow. Neither food nor water were withheld. Drinking water was supplemented with prophylactic antibiotics (1.2 mM Neomycin trisulfate and 0.13 mM Polymyxin B sulfate). To minimize non-hematopoietic toxicity, a non-lethal dose of 200 cGy was administered to NSG mice. BM cells were replenished with donor cells injected within 4 hours after irradiation. For retroorbital injection of HSPCs, the mouse was anesthetized using inhaled isoflurane at 4% for induction and 1.5-2% during the procedure, delivered in 100% oxygen at a rate of 1.5-2 L/minute. The mouse was kept warm on a warming pad. A sterile 27-gauge needle attached to a sterile 1.0 ml syringe was used for the injection of CD34+ cells. After a surgical plane of anesthesia had been achieved, the needle was inserted between the medial canthus and the globe. The tip of the needle was directed at the rostrodorsal aspect of the orbit and gently advanced toward the ophthalmic plexus. A volume of up to 200 μl was injected. When the injection was completed, the needle was removed, and the mouse was observed for bleeding at the injection site. For blood collection, the mouse was manually restrained in the operator’s non-dominant hand by securing the loose skin on the dorsal head, neck, and thorax with the thumb and index finger. The puncture site was identified 2 mm from the “dimple” near the oral commissure, along an imaginary line from the dimple to the pinna. A lancet of appropriate size is used to puncture the skin and vein. Blood was allowed to drip into a collection vial filled with 5 ml EDTA-PBS solution. When bleeding has stopped, manual restraint was released, providing hemostasis. The mouse was placed back in the cage and observed for additional bleeding as well as for signs of respiratory distress. Blood collection was performed once a month.

### Flow cytometry

RBC-free cell pellets were processed and cryopreserved in 4-5 aliquots as described ^35,37^. Cryopreserved cells, including tibial FBM and PBMCs, were analyzed by flow cytometry using monoclonal antibodies validated for binding to rhesus macaque proteins ^38^ and lymphoid-specific and myeloid-specific gating strategies. For FBM phenotyping, 5×10^5^-1×10^6^ cells were stained with antibodies to CD45 (D058-1283, Biolegend, San Diego, CSA, USA), CD3 (clone SP34, BD Pharmingen), CD20 (clone 2H7, Biolegend), CD8 (clone SK1, Biolegend), CD14 (clone M5E2, Biolegend), CD16 (clone 3G8, Biolegend), HLA-DR (clone L243, Biolegend), CD11b (clone ICRF44, Biolegend), CD163 (clone GHI/61, Biolegend), and Mac387 (AB7429, Abcam, Cambridge, England) and analyzed on an LSRII flow cytometer (BD Biosciences, San Jose, CA). For blood phenotyping, 5×10^5^ freshly thawed PBMCs were stained with CD3 (clone SP34, BD Biosciences, San Jose, CA), CD8b (clone 2ST8.5H7, Beckman Coulter), CD20 (clone 2H7, Biolegend), CD14 (clone M5E2, Biolegend), HLA-DR (clone L243, Biolegend), CD11c (clone 3.9, Biolegend), and CD16 (clone 3G8, Biolegend). Cells were incubated at 4C for 30 minutes, washed twice in FACS buffer (1X PBS with 2% FBS and 1 mM EDTA) and the data were acquired using the Attune NxT flow cytometer (ThermoFisher). For FL phenotyping, 3×10^5^ cryopreserved, viable liver mononuclear cells were thawed and stained with antibodies to CD3 (clone SP34, BD Pharmingen), CD20 (clone 2H7, Biolegend), CD8b (clone 2H7, Thermo), CD4 (clone L200, BD Pharmingen), CD14 (clone TUK4, Miltenyi Biotec), CD11b (clone ICRF44, Biolegend), for 20 minutes at 4C with anti-human Fc Block (Miltenyi Biotec). Cells were incubated with DAPI viability dye (Thermo) for the last minute of the incubation. Cells were then passed through a 100 μm cell strainer (Corning) into FACS buffer (1X PBS with 2% FBS and 1 mM EDTA) and analyzed with a FACS Aria flow cytometer (BD Biosciences).

### TLR stimulation assays

FBM cells (5×10^5^) were stimulated with LPS (TLR2/4 agonist 1 μg/ml), Pam3CSK4 (TLR1/2 agonist, 1 μg/ml) or HKLM (TLR2 agonist, 10^8^ particles/ml) for 8 hours in the presence of 5 μg/ml brefeldin A (Sigma-Aldrich). TLR agonists were purchased from InvivoGen (San Diego, CA, USA). Cells were stained with antibodies to CD11b, HLA-DR, and CD163 to delineate monocytes/macrophages. Following surface staining, cells were permeabilized and stained with monoclonal antibodies to TNFα (clone MAb11, Invitrogen). Samples were washed in FACS buffer and analyzed on an LSRII flow cytometer. Data were analyzed using FlowJo v10 (Tree Star, Ashland, OR, USA) and Prism v6 (GraphPad Software, San Diego, CA, USA).

### FACS sorting and scRNA-Sequencing of fetal CD34+ cells

Freshly thawed antibody-labeled CD34+ HSPCs were enriched for live cells by FACS using SYTOX Blue dead cell stain (ThermoFisher) on a BD FACS Aria Fusion cell sorter. Cells were sorted into RPMI containing 30% FBS, 5% glutamine, and antibiotics. Cells were then counted in triplicates on a TC20 Automated Cell Counter (Bio-Rad), washed and resuspended in 1X PBS with 0.04% BSA to a final concentration of 1200 cells/μL. Single cell suspensions were then immediately loaded on a 10X Genomics Chromium Controller with a loading target of 17,600 cells. Libraries were generated using the 3’ V3 chemistry per the manufacturer’s instructions (10X Genomics, Pleasanton, CA, USA) and sequenced on an Illumina NovaSeq with a sequencing target of 30,000 reads per cell.

### Single cell RNA-Seq data analysis

Raw reads were aligned and quantified using the Cell Ranger’s *count* function (version 4.0, 10X Genomics) against the *Macaca mulatta* reference genome (MMul10) using the STAR aligner. Downstream processing of aligned reads was performed using Seurat (version 3.2.2). Droplets with ambient RNA (cells with fewer than 400 detected genes), potential doublets (cells with more than 4000 detected genes) and dying cells (cells with more than 20% total mitochondrial gene expression) were excluded during the initial quality control. To rule out biases in different stages of cell cycle, cells were then scored for S or G2/M phase using a catalog of canonical markers ^39^. Cells with high ribosomal gene expression (≥ 30 pc of total gene expression) were further excluded before integration. Data objects from the two groups were integrated using Seurat’s *IntegrateData* function ^40^. Data normalization and variance stabilization were performed using *SCTransform* function using a regularized negative binomial regression to correct for differential effects of mitochondrial and ribosomal gene expression profiles. Dimension reduction was performed using the *RunPCA* function to obtain the first 30 principal components followed by clustering using the *FindClusters* function in Seurat. Clusters were visualized using UMAP algorithm as implemented by Seurat’s *runUMAP* function. Cell types were assigned to individual clusters using *FindMarkers* function with a fold-change cutoff of at least 0.4 and using a known catalog of well-characterized scRNA markers for HSCs (**Supplementary Table S5**) ^41^.

### Pseudo-temporal and differential expression analyses

Hematopoietic lineages and developmental pseudo time of myeloid cells was inferred by Slingshot. The annotated clusters inferred by KNN graph were used as input for Slingshot. Differential expression analysis of the chow and WSD groups within each cell type was performed using the non-parametric Wilcoxon rank sum test in Seurat. Genes with at least a 0.4-fold change (log_2_ scale) and with an adjusted p-value ≤ 0.05 were considered significant. Pathway analysis of differential signatures was performed using Enrichr. For module scoring, we compared gene signatures and pathways from KEGG (https://www.genome.jp/kegg/pathway.html) in subpopulations using Seurat’s *AddModuleScore* function. Group differences were compared using unpaired t-test followed by Welch’s correction.

### Statistical analyses

All statistical analyses were conducted in Prism 8 (GraphPad). A correlation analysis between maternal and petal parameters was conducted by calculating the Pearson correlation coefficients and their associated p-values. The p-values were then adjusted using Benjamini-Hochberg procedure to control the false discovery rate (FDR) below 0.05 for selected maternal fetal parameters. This method was implemented in R using the “p.adjust” function. All two-group comparisons were tested for statistical differences using unpaired t-tests.

## RESULTS

### Maternal WSD induces fetal bone marrow adipogenesis during the 3^rd^ trimester

Animals consuming the WSD were heavier and exhibited a significantly greater percentage of total body fat than control animals ^34,42^. WSD-fed animals also remained heavier before and during pregnancy than control animals (**Supplementary Figure S1B and C**). While fetal weights were not significantly affected by maternal diet, fetal retroperitoneal fat pad weights and maternal and umbilical artery glucose levels were significantly elevated in the WSD group (**Supplementary Figures S1D-G**). During the 3rd trimester, the migration of fetal HSPCs to the FMB is temporarily linked to bone ossification and formation of the BM medullary cavity in long bones (**Figure 1A**). To elucidate the effects of maternal WSD on the development of fetal bones and the FBM niche, we performed micro-Computed Tomography (microCT) and histological evaluation of fetal femurs. As expected for immature bones, the femurs of control and WSD fetuses were highly trabeculated, but the latter exhibited more open central medullary cavities in bone shafts compared to controls (**Figure 1B and C**). While the average trabecular thickness in fetal femurs was not significantly different between the WSD and control groups, the trabecular thickness in distal femur was positively correlated with maternal glucose levels at C-section (**Figure 1E and Supplementary Table S1**). Additionally, the number of trabeculae per bone volume (trabecular density) was significantly reduced in the distal femurs of WSD fetuses (**Figure 1F).**

**Figure 1.**
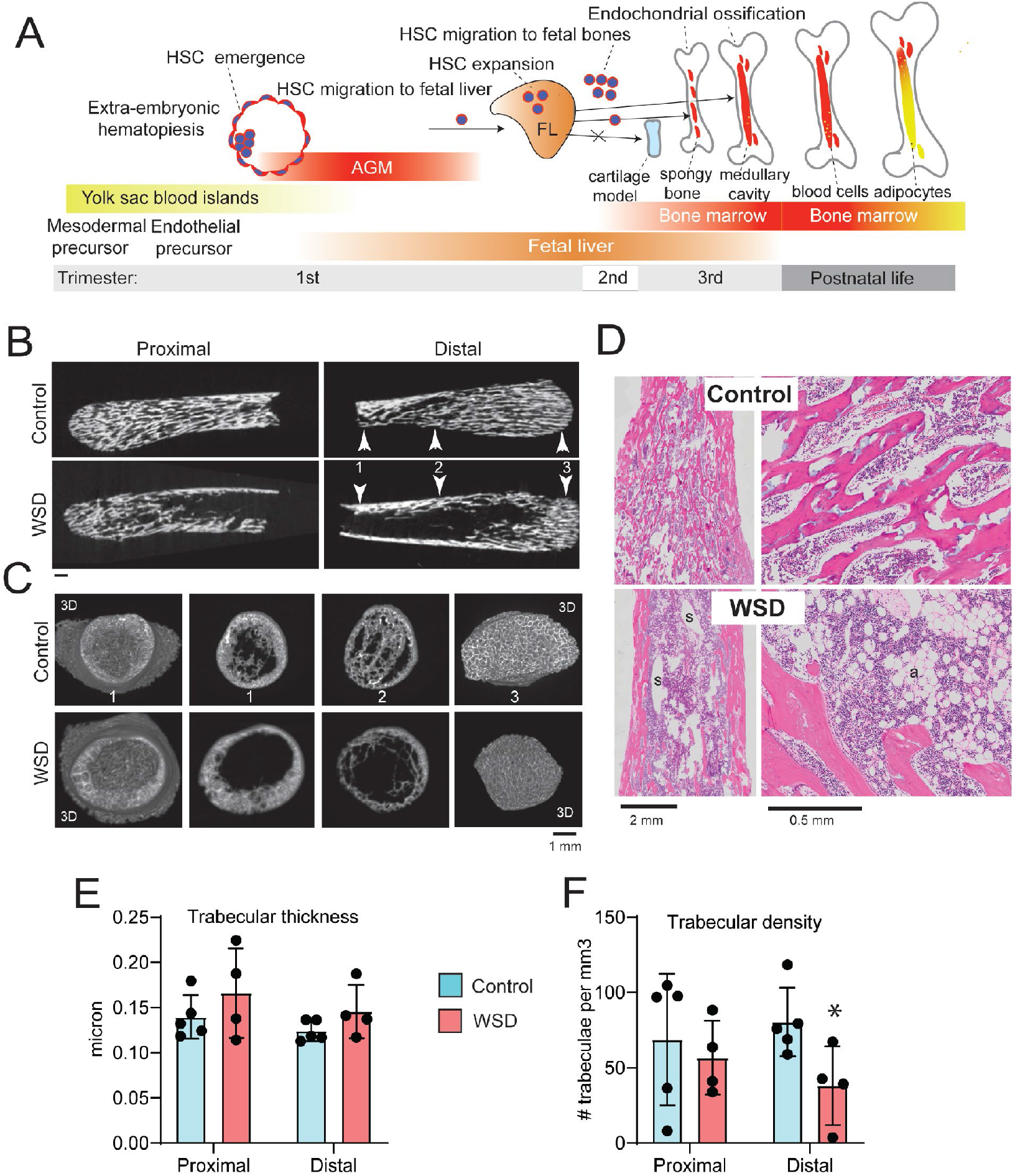
Maternal WSD impacts fetal bone development and BM adipogenesis. (A) Migration of HSPCs between hematopoietic sites during development. Human HSPCs emerge in the aorta-gonad-mesonephros (AGM) during the first trimester (E26-40) and FL colonization starts at E28. During the 3rd trimester, HSPCs migrate from the FL to the FBM. Fetal bones are highly trabeculated. Bone marrow adipogenesis begins during postnatal life and is associated with the formation of the central medullary cavity. (B) Representative longitudinal midline cross-sections across the bone shafts of the proximal and distal femurs of control and WSD fetuses. Arrowheads and numbers point at the location of radial cross-sections shown in panel “C.” (C) Radial cross-sections across the distal femurs. Far-left and far-right images represent the 3D view of the femurs, looking from an open and closed ends, respectively. Middle panels (positions 1 and 2) represent the view of the 1-mm thick slices across fetal femurs. (D) Representative images of H&E stained proximal femurs from the control and WSD fetuses; a, adipocytes; s, sinusoids. (E) Mean trabecular thickness and (F) connectivity density across the proximal and distal femur. Bars are means ± SEM; t-tests; *p<0.05.

The BM cavities of fetal femurs derived from the WSD group were densely populated by adipocytes, while the femurs of control fetuses only occasionally contained adipocytes (**Figure 1D**). Because quantitative immunohistochemical analysis across the fetal femur was difficult to perform due to large size and extensive trabeculation of fetal bones, we performed additional analysis of fetal sterna, representing an alternative site of hematopoiesis. Maternal WSD induced a significant increase in the number of adipocytes in fetal sterna, while adipocytes were only occasionally detected in control sterna (**Supplementary Figures S2A and B**). Furthermore, the sternal trabeculae from control fetuses contained a higher proportion of cartilage tissue compared to WSD fetuses, whose trabeculae were completely ossified (**Supplementary Figures S2A, yellow arrowheads**). We next analyzed whether maternal WSD exposure changes the cellular composition and anatomical localization of macrophages and HSPCs in fetal sterna. Both cell types were distributed throughout the FBM parenchyma (**Supplementary Figures S2C, F and I**). The total number of CD68+CD163+ macrophages and CD34+ HSPCs per sternal section were not significantly different between the control and WSD groups (**Supplementary Figures S2D and G**). However, a significant higher proportion of macrophages and HSPCs was found in close proximity to FBM adipocytes in the sterna of WSD fetuses compared to controls (**Supplementary Figures S2E and H**). These data demonstrate that maternal WSD induces premature FBM adipogenesis in fetal bones.

### Characterization of rhesus macaque fetal HSPCs at the single-cell resolution

We hypothesized that the WSD-induced changes in the FBM niche impact the differentiation and functional properties of fetal HSPCs. First, we determined the transcriptional landscape of FBM HSPCs at single-cell resolution. We developed a bone crushing method that allowed us to obtain the sufficient number of HSPCs from fetal long bones. HSPCs were isolated from control and WSD-exposed fetuses (n=3/group) using NHP-specific anti-CD34 antibodies, before being profiled by droplet-based scRNA-Seq (**Supplementary Figure S1A and S3A**). Dimension reduction and clustering of CD34+ single-cell RNA profiles revealed that transcriptomes of HSPCs from control and WSD-exposed fetuses were largely overlapping (**Supplementary Figure S3B**), with a total of 21 cell clusters identified (**Supplementary Table S2**). Consolidation of redundant clusters resulted in fewer hematopoietic progenitor clusters organized in a continuum state (**Figure 2A**). These clusters were annotated based on markers highly expressed in each cluster relative to the remaining clusters (**Supplementary Table S2**). This approach allowed us to identify a cluster of hematopoietic stem cells (HSCs) expressing high levels of *HOXA9 and HOPX*, proliferating HSCs marked by high *MKI67* ^43^), megakaryocytic-erythroid progenitors (MEPs) expressing *KLF1* and *HBA*, common lymphoid progenitors (CLPs) expressing *IL7R*, *RAG1* and *DNTT*, pre-conventional dendritic cell precursors (cDCs) expressing *CCR2*, *MAMU-DRA* and *CD74*; pre-plasmacytoid DCs (pDCs) expressing *IRF7* and *CCL3*, granulocyte-myeloid progenitors (GMPs) expressing *ELANE* and *MPO*; common monocyte progenitors (cMoPs) expressing *OAZ1* and *LYZ*, and pro-monocytes expressing myeloid cell markers *S100A8*, *IL1B* and *VCAN* ^44–47^ (**Figure 2B-D; Supplementary Figure S4; Supplementary Table S2**).

**Figure 2.**
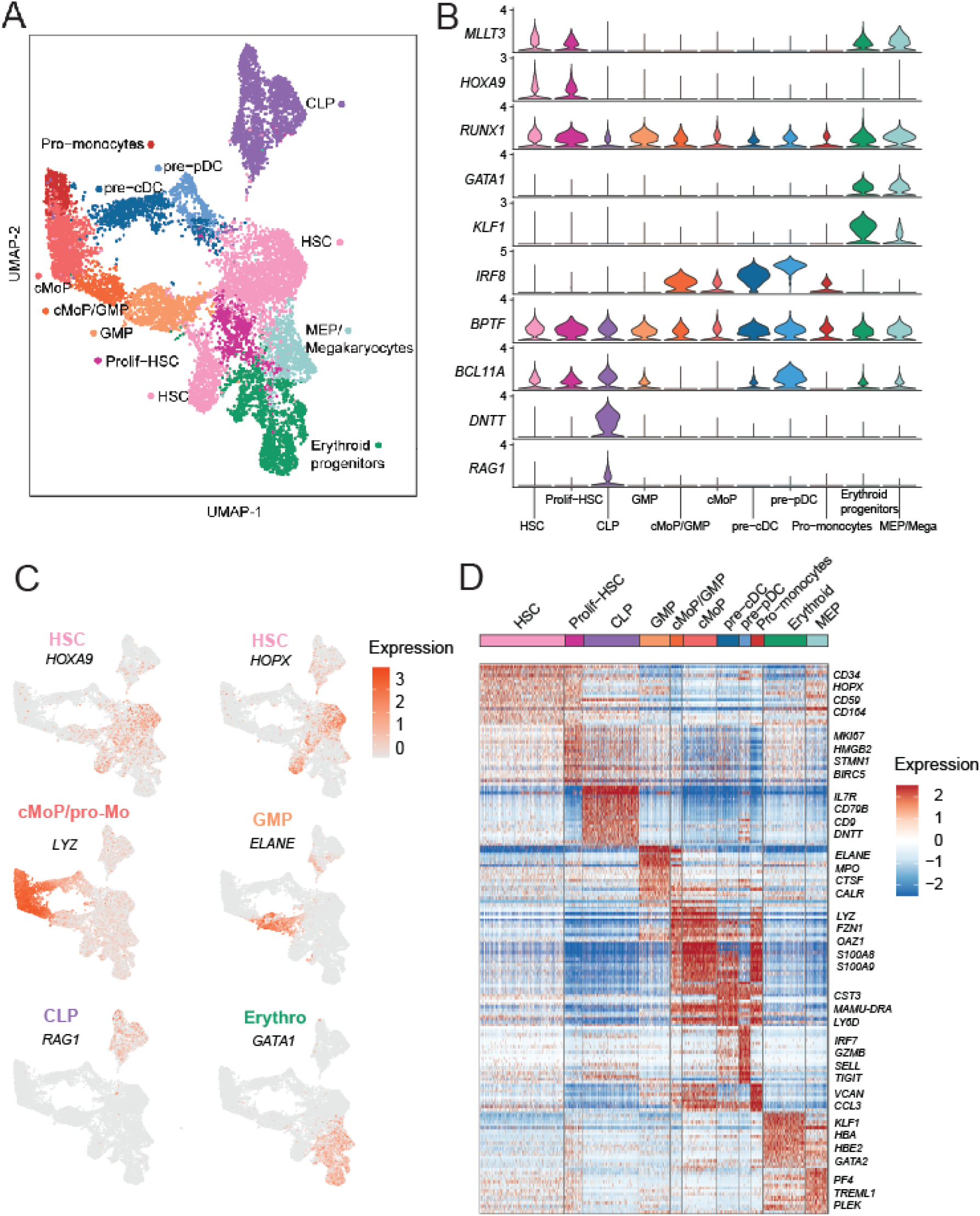
Single-cell transcriptome of fetal macaque bone marrow HSPCs. (A) Uniform Manifold Approximation and Projection (UMAP) representation of single-cell clusters after dimension reduction of sorted CD34+ HSPCs (n=3/group, pooled). Progenitor names are indicated. (B) Expression levels of transcriptional regulators of HSPC differentiation identified in the present study. (C) Expression levels of key lineage-specific markers (gene names are shown in black) in HSPCs; progenitor names are color-coded. (D) Heat map of progenitor-specific genes; only a subset of lineage-specific genes is shown.

Furthermore, as observed in fetal human HSCs, fetal macaque HSCs showed high expression levels of the transcription factor gene *MLLT3*, the master regulator of human HSC maintenance ^48^. Additionally, fetal rhesus macaque HSCs expressed high levels of *CD164* encoding the sialomucin adhesive glycoprotein expressed in the most primitive branches of human CD34+ HSPCs ^41^. Fetal macaque HSCs also expressed *HOXA9,* the master transcriptional regulator of HSC differentiation in humans ^49,50^, as well as *RUNX1* encoding the essential transcription factor required for HSC maintenance ^51^ (**Figures 2B-D; Supplementary Figure S4**). We also confirmed that rhesus macaque and human ^52^ fetal HSPCs share lineage-specific genes expressed in megakaryocyte-erythroid progenitors (*GATA1/2, PLEK, KLF1, MS4A2,* and *BLVRB*), granulocyte-monocyte progenitors (*AZU1, LYZ,* and *MPO*), DC progenitors (*IRF*8), and lymphoid progenitors (*HMGB2* and *CD79B*) (**Figures 2B-D; Supplementary Figure S4; Supplementary Table S2**). *CD38* was broadly expressed in fetal HSPCs (**Supplementary Figure S4**), consistent with a previous study, suggesting an overlap in transcriptional states between CD34+CD38- and CD34+CD38+ human HSPCs ^41^. Furthermore, CLPs formed a distinct cell cluster and exhibited a unique gene expression pattern characterized by high expression of *DNTT* and *RAG1* (**Figures 2B and C, Supplementary Figure S4; Supplementary Table S2**). The CLP cluster was further sub-divided into CLP-1 (expressing the T-cell marker *CD3D*), CLP-2 (expressing the B-cell markers *CD79B* and *MAMU-DRA*), and proliferating CLPs (expressing high levels of *MKI67*) (**Figures 4C and Supplementary Figure S4**). These data demonstrate that the single-cell transcriptome of rhesus macaque FBM HSPCs exhibits high dimensionality and significant similarities with human fetal HSPCs.

**Figure 3.**
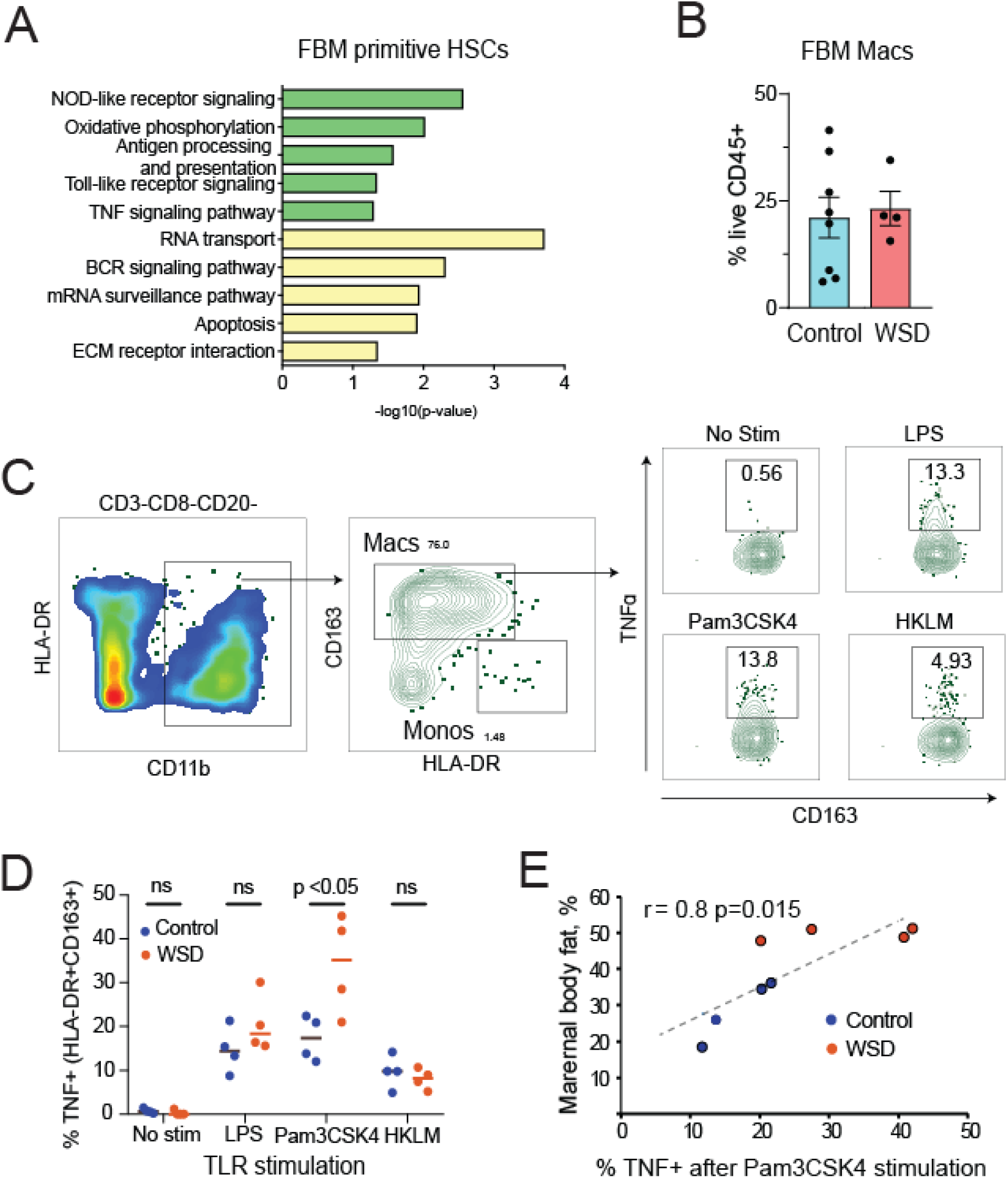
Maternal WSD induces a hyperinflammatory phenotype in FBM progenitors and macrophages. (A) Pathway analysis of differentially expressed genes in primitive HSCs in response to a maternal WSD; green, upregulated genes; yellow, down-regulated genes. (B) Macrophage frequencies in the tibial FBM of control and WSD fetuses; each data point represents one fetus; bars are means ± SEM. (C) Gating strategy used for studying toll-like receptor ligand responses in FBM macrophages. (D) TNFα responses in FBM macrophages following TLR ligand-stimulation; each data point shows the % of TNFα+ cells within the CD11b+HLA-DR+CD163+ gate; n=4 control fetuses and 4 WSD fetuses; horizontal bars are means. All two-group comparisons were tested for statistical differences using unpaired t-test. (E) Correlation between maternal body fat content before pregnancy and macrophage responses to Pam3CSK4 stimulation; Pearson coefficient (r) and p-values are indicated.

**Figure 4.**
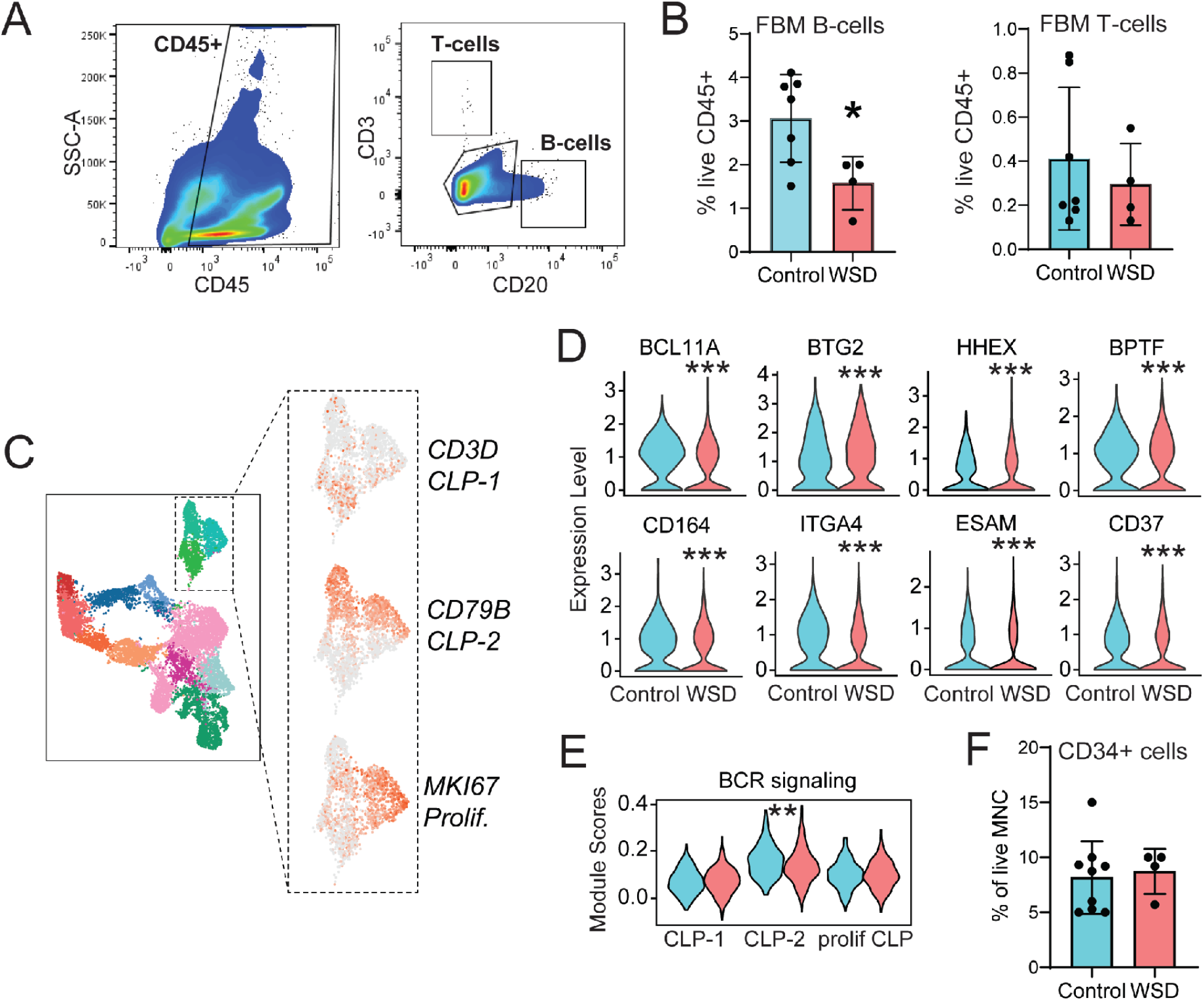
Maternal WSD attenuates fetal B-cell development in long bones. (A) The expression of lineage-specific transcripts in CLPs. The spatial positions and expression of specific genes in CLP subtypes are indicated. (B) Normalized RNA levels of transcriptional regulators (upper row) and adhesion molecules (bottom row) downregulated (p<0.001) in fetal CLPs with a maternal WSD. (C) Violin plot comparing module scores for B-cell receptor (BCR) signaling within the CLP clusters. (D) Fetal CD34+ cell frequencies in FBM; CD34+ cells were isolated from the tibial mononuclear cell (MNC) fraction using the anti-CD34 antibody. Each data point represents a % of CD34+ cells within live FBM MNCs. (E) Gating strategy used for identifying FBM CD3+ T-cells and CD20+ B-cells within the parental CD45+ gate. (F) B-cell and T-cell frequencies in the CD34-CD45+ fraction of the FBM. Each data point represents one fetus. Bars are means ± SEM. All two-group comparisons were tested for statistical differences using unpaired t-tests; *p<0.05; **p<0.01; ***p<0.001.

### Maternal WSD induces a proinflammatory phenotype in fetal hematopoietic progenitors and bone marrow macrophages

The analysis of differentially expressed genes (DEGs) revealed that maternal WSD exposure leads to upregulation of pathways associated with myeloid cell activation, including NOD-like receptor (NLR), toll-like receptor (TLR), and tumor necrosis factor (TNF)α signaling pathways in in FBM HSCs (**Figures 3A**). Moreover, the proinflammatory genes *S100A8/9*, *NFKB1A*, and *PTGS2* were highly upregulated in both HSCs and cMoPs (**Supplementary Table S3**). We next determined if this transcriptional signature impacts the composition and functional properties of myeloid cells in fetal long bones. As described for sternal macrophages, long-bone macrophage frequencies were not affected by maternal WSD (**Figures 3B**). However, FBM macrophages from the WSD group produced significantly higher levels of TNFα than those from the control group following stimulation with a synthetic triacylated lipopeptide Pam3CysSerLys4 (Pam3CSK4, a TLR1/2 ligand) (**Figure 3C and D**). Moreover, the magnitude of the TNFα response to Pam3CSK4 by FBM macrophages positively correlated with maternal body fat measured prior to pregnancy (**Figure 3E**). The frequencies of myeloid subsets residing in fetal blood and the FL were not affected by a maternal WSD exposure (**Supplementary Figure S5**). Collectively, these data demonstrate that a maternal WSD exposure results in heightened proinflammatory responses by FBM macrophages but had no effect on FBM myeloid cell frequencies.

### Maternal WSD attenuates B-cell development in fetal bones

After investigating the effect of maternal WSD on fetal myelopoiesis, we assessed its effect on lymphopoiesis. Flow cytometry analysis of CD34-CD45+ immune cells showed that maternal WSD reduced B-cell content in the FBM. In contrast, proportions of FBM CD3+ T- lymphocytes that mature in the thymus and can subsequently be recruited into the BM ^53^ were comparable between the two groups (**Figure 4A and B**). The frequencies of lymphoid subsets residing in fetal blood and FL were not affected by maternal WSD (**Supplementary Figure S5**). We next explored whether a maternal WSD alters B-cell development in lymphoid progenitors at the transcriptome level. The analysis of DEGs in fetal CLPs and HSCs revealed a significant decrease in expression of several transcriptional regulators (*BCL11A*, *BTG2, HHEX*, *BPTF*, *NCL* and *MLLT3*) and cell adhesion molecules (*CD164, ITGA4, ESAM and CD37*) in WSD-exposed fetuses (**Figure 4C and D; Supplementary Table S3**). The *MLLT3* ^48^ and *CD164* ^41^ genes have been previously shown to be essential for the maintenance and the hematopoietic reconstitution ability of HSPCs. The most prominent downregulated genes included *BTK,* required for pre-B-cell differentiation ^54^, *ERG,* which is essential for lymphopoiesis ^49,55,56^, as well as genes involved in the lymphoid differentiation pathway (*ETV6*, *SMARCB1*, *GRB2*, *SYK*, and *TXNDC12*) (**Supplementary Table S3**). Furthermore, we detected significant downregulation of *BCL11A,* required for fetal hematopoiesis ^57^; *IRF3*, the lymphoid transcription factor involved in the interferon signaling pathway ^58^; *CD164* and *ITGA4* (which were also downregulated in HSCs); as well as *CD79A, VPREB1, RAG1*, and *HMGB2* which play a critical role in lymphoid lineage development in human fetuses ^52^ (**Figure 4D and Supplementary Table S3**). The DEG analysis revealed a suppression of B-cell receptor signaling in B cell progenitors (CLP-2) with maternal WSD (**Figure 4C and E**). However, the proportions of CD34+ HSPCs within the mononuclear FBM fraction were not significantly different between the two groups (**Figure 4F**). Collectively, our analyses indicate that a maternal WSD changes the transcriptional program of FBM CLPs, potentially dysregulating B-cell development.

### Maternal WSD alters the engraftment ability of FBM HSPCs under regenerative stress

We next tested whether maternal WSD alters HSPC function in vivo. We transplanted 120,000 tibial FBM CD34+ HSPCs into nonlethally irradiated immunodeficient NOD/SCID/IL2rγ- /- (NSG) mice that were maintained on a low-fat chow diet throughout the engraftment period (**Supplementary Figure S6A and B**). Thirteen weeks following engraftment, the levels of macaque CD34+ cells in the BM were significantly lower in the mice engrafted with HSPCs from the WSD group compared to those engrafted with control HSPCs (**Figure 5A and B**). We observed no significant group differences in frequencies of B and T-cells in the BM and blood of NSG mice (**Figure 5C and D**). In contrast, frequencies of myeloid cells, including monocytes, and granulocytes (gating strategy developed by ^27^) were significantly reduced in the BM of NSG mice that received CD34+ cells from maternal WSD fetuses (**Figures 5E, F and G**). To test whether a maternal WSD alters the intrinsic differentiation properties of FBM HSPCs, we isolated engrafted CD34+ cells from the BM of NSG mice and conducted a colony-forming assay (CFU) using 250 CD34+ cells, derived from the control and WSD groups. There were no significant group differences in the number of granulocyte–macrophage progenitor colony forming units (CFU-GM) or erythroid burst forming units (BFU-E) (**Supplementary Figure S6C**), suggesting that control and WSD HSPCs exhibit similar in vitro erythro-myeloid differentiation abilities. In summary, our results demonstrate that a maternal WSD impacts the intrinsic properties of FBM HSPCs, thus affecting their engraftment in the BM under regenerative conditions.

**Figure 5.**
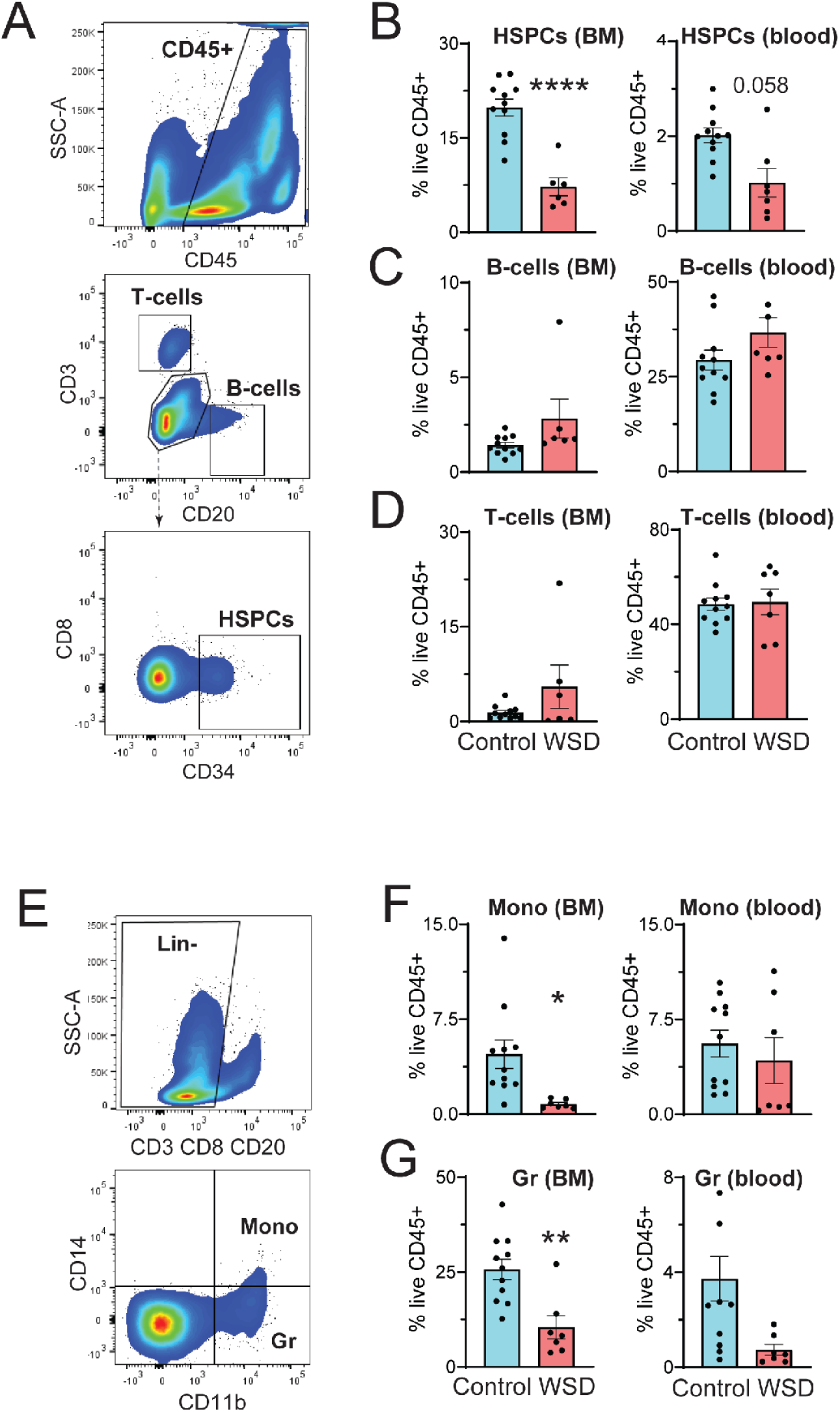
Maternal WSD impairs the engraftment ability of FBM HSPCs. (A) Outline of engraftment experiments (see “Materials and Methods” for details). (B) Gating strategy used for the analysis of rhesus CD3+ T-cells and CD20+ B-cells within the CD45+ gate in NSG mice. Rhesus macaque CD34+ HSPCs are identified within the CD45+CD3-CD20-gate. (C) HSPC, (D) B-cell, and (E) T-cell frequencies in the CD45+ fraction of the BM and blood of NSG mice. (F) Gating strategy used for identifying myeloid cells. Rhesus macaque monocytes (Mono) and granulocytes (Gr) are identified within the Lin-gate as CD14+CD11b+ or CD14-CD11b+ cells, respectively. (G) Monocyte and granulocyte frequencies in the BM and blood of NSG mice. Each data point represents one NSG mouse; bars are means ± SEM; t-test, *p<0.05, **p<0.01, ****p<0.0001.

## DISCUSSION

Maternal obesity adversely impacts immune function in offspring; however the mechanisms remain largely unknown. Experimental and clinical obesity in adults has been linked to the dysregulation of hematopoiesis ^59–61^ mediated in part via gut-derived microbial-derived metabolites and TLR ligands ^62^. Previous studies have also shown that maternal WSD compromises FL hematopoiesis ^8^, but very little is known about the effect of WSD on FBM hematopoiesis during fetal life in models that resemble human development. Here we show for the first time that maternal WSD increases FBM adiposity during development in utero, and also demonstrate that NHP hematopoietic markers share lineage-specific genes closely aligned with those of humans. While hematopoietic precursor pools were not changed, we found that maternal WSD shifts fetal gene expression patterns in FBM HSPCs and macrophages significantly toward inflammation. This shift significantly compromised the engraftment ability of FBM HSPCs under regenerative conditions. Overall, our findings suggests for the first time that maternal WSD affects FBM architecture and adiposity, while increasing the inflammatory potential of HSPCs and FBM macrophages, with significant perturbations in genes critical for B-cell development at the single-cell level.

Focusing on FBM, we found that maternal WSD exposure impacts bone development and triggers premature FBM adipogenesis. The presence of adipocytes in the BM cavity is associated with a postnatal state, as reported by previous NHP ^35^, rodent, and human ^63,64^ studies. The pattern of fetal BM adipogenesis induced by a maternal WSD also resembles age-related changes in adult long bones associated with the replacement of hematopoietic marrow with adipogenic marrow ^63,65–67^. In addition, we found that bones from the WSD fetuses are more ossified compared to control fetuses. It is known that the medullary cavity of long bones in human and rodent fetuses is composed of partially ossified trabeculae comprised of woven bone, with BM spaces interspersed between the bony trabeculae ^68^. This structural organization of fetal bones is consistent with the bone phenotype observed in macaque fetuses, with WSD fetuses showing more developed trabeculae-free medially cavities than control fetuses. These findings are in line with previous observations showing that a high-fat diet increases expansion of the adipogenic, but not the osteochondrogenic lineage in adult mice ^65^. BM adipocytes have been initially described as the negative regulators of hematopoiesis under regenerative stress in mice^69^. These findings are also consistent with our transplantation data showing that a maternal WSD has a negative impact on HSPC engraftment in NSG mice.

We also demonstrated that maternal WSD evokes a persistent inflammatory gene signature in FBM HSPCs and a latent hyperinflammatory phenotype in FBM macrophages. These findings suggest that a maternal WSD in utero regulates fetal immune system development, in part, through reprogramming fetal HSPC function. This study, therefore, is the first demonstration in NHPs that maternal diet can modulate developmental hematopoiesis in the BM in utero. Given the negative effects of maternal WSD on hematopoietic function in long bones in NHP fetuses, alongside increased adipocyte content, we speculate that excess BM adiposity in utero has a negative effect on specific aspects of hematopoiesis. However, it is possible that other components of the FBM niche (such as stromal and bone cells) are also affected by a maternal WSD, and more work is necessary to understand how alterations in the cross-talk between adipocyte-rich microenvironment and HSPCs affect the subsequent development and function of the immune system. For example, we have previously shown that adult macaque BM adipose tissue releases factors enriched in extracellular vesicles ^35^, representing the potential regulators of HSPC function and differentiation ^70–72^. Likewise, there may be maternal metabolic signals from the maternal microbiome that play a role in prenatal dietary priming of hematopoietic progenitors and contribute to developmental programming ^73^.

Our immunophenotypic analysis shows that a maternal WSD also reduced B-cell numbers in fetal long bones. Focusing on FBM CD34+ HSPCs, our scRNA-Seq analysis revealed that a maternal WSD reduced the expression of B-cell development genes, including *VPREB1* ^74^, *VPREB3* ^75^, *BST*2 ^76^, and *CD79a* ^77^ in FBM CLPs. We also demonstrate that FBM HSPCs from the WSD group exhibited reduced engraftment and myeloid lineage reconstitution abilities in nonobese recipient mice, which are consistent with our scRNA-Seq data showing that maternal WSD results in downregulation of cell adhesion molecules *CD164, ITGA4, ESAM and CD37* in FBM HSPCs. These data are consistent with the concept that maternal WSD drives intrinsic changes in HSPCs that affect their biological function in the FBM. We did not observe significant effects of maternal WSD on myeloid cell numbers in fetal bones, FL, and peripheral blood. In addition, a previous study in mice has shown that in the FL, maternal obesity together with a maternal high-fat diet, skewed HSPCs toward myeloid-biased differentiation which was niche-dependent ^8^. This suggests that chronic high-fat diet may bias FL HSPCs toward myeloid differentiation at the expense of HSC self-renewal. Furthermore, our studies showed that WSD enhanced TLR1/2 cytokine responses by FBM macrophages, which is consistent with rodent studies that demonstrated hyper-inflammatory responses in BM-derived macrophages ^78^ and microglia ^79^ of offspring born to dams on a high-fat diet and with our scRNA-Seq analysis showing the upregulation of TLR and TNFα signaling pathways and proinflammatory genes *S100A8/9*, *NFKB1A*, and *PTGS2* in monocyte progenitors from the WSD fetuses. Our study therefore expands previous reports that obesity drives peripheral adipose tissue inflammation ^80–83^ and suggests that diet-induced expansion of BM adipose tissue is also associated with an increase in a proinflammatory environment in the FBM.

In summary, this study demonstrates that maternal WSD exposure results in altered fetal hematopoiesis. Specifically, maternal WSD led to (1) an increase in BM adipogenesis in fetal bones; (2) enhanced trabeculae ossification in fetal bones; (3) an increase the formation of medullary cavities in fetal bones; (4) an increase in proinflammatory cytokine production by FBM macrophages; (5) a decrease in B-cell development in long bones; and (6) a reduction in the regenerative abilities of HSPCs following transplantation in recipient mice. These changes in the FBM resemble an aging-like phenotype, which is also associated with an increase in BM adiposity, reduced lymphopoiesis, an increase in inflammation, and reduced HSPC function^64,84,85^. Future studies are needed to understand how maternal obesity affects the interactions between HSPCs and various types of the BM microenvironments. These results set the stage for understanding the link between maternal obesity, prenatal nutrition, and diseases involving immune progeny of the HSPC compartment in children ^24,86,87^, and highlight the need to better understand the susceptibility of the developing hematopoietic system to metabolic dysregulation effects across the lifespan.

## AUTHOR CONTRIBUTIONS

scRNA-Seq assay and bioinformatic analysis were generated by SS. Immunohistochemistry was conducted by CNC and LP. HSPC transplantation and cell isolation in NSG mice were performed by OV, JJR, and DG. Immunophenotyping experiments were performed by BJB, SS, OV, MJN, and JJR. Flow cytometry data were analyzed by IM, OV, SS, DG, MJN, EMP, and BJB. Fetal tissue collection was performed by DT, JJR and OV. Statistical analysis was performed by OV, SS, BJB, and WS. MicroCT analysis was done by JRL and WP. The manuscript was written by OV, SS, and IM. OV, JDH, MJN, JEF, SRW, and CTR contributed to the conception of, provided access to samples, and edited the manuscript.

## ACKNOWLEDGEMENTS

We thank Dr. Lauren Drew Martin for help with blood collections. We thank the head of the Small Lab Animal Unit, Kati Marshall, for help in animal handling and transplantation. We thank Dr. Jennifer Atwood (UCI Flow Cytometry Core) for assistance with cell sorting and Dr. Melanie Oakes (UCI GHTF) for assistance with library preparation and sequencing of 10x samples. This study was supported by NIH grants P50 HD071836 to CTR and JDH, P51 OD01192 for operation of the Oregon National Primate Research Center, 1R01AI142841 and 1R01AI145910 to IM, R24-DK090964 to JEF, F30-DK122672 to MJN, R01-DK108910 to SRW, and R01-DK119394 to EMP.

## DATA SHARING

Gene expression data are available at the NCBI SRA data repository under Accession Number PRJNA723061.

## REFERENCES

1. Lashen H, Fear K, Sturdee DW. Obesity is associated with increased risk of first trimester and recurrent miscarriage: matched case-control study. Hum Reprod 2004; 19(7): 1644–6.

2. Stothard DJ, Hopkins JM, Burns D, Dunn MH. Stable, continuous-wave, intracavity, optical parametric oscillator pumped by a semiconductor disk laser (VECSEL). Opt Express 2009; 17(13): 10648–58.

3. Griffiths PS, Walton C, Samsell L, Perez MK, Piedimonte G. Maternal high-fat hypercaloric diet during pregnancy results in persistent metabolic and respiratory abnormalities in offspring. Pediatr Res 2016; 79(2): 278–86.

4. Rastogi S, Rojas M, Rastogi D, Haberman S. Neonatal morbidities among full-term infants born to obese mothers. J Matern Fetal Neonatal Med 2015; 28(7): 829–35.

5. Suk D, Kwak T, Khawar N, et al. Increasing maternal body mass index during pregnancy increases neonatal intensive care unit admission in near and full-term infants. J Matern Fetal Neonatal Med 2016; 29(20): 3249–53.

6. Odaka Y, Nakano M, Tanaka T, et al. The influence of a high-fat dietary environment in the fetal period on postnatal metabolic and immune function. Obesity (Silver Spring) 2010; 18(9): 1688–94.

7. Myles IA, Fontecilla NM, Janelsins BM, Vithayathil PJ, Segre JA, Datta SK. Parental dietary fat intake alters offspring microbiome and immunity. J Immunol 2013; 191(6): 3200–9.

8. Kamimae-Lanning AN, Krasnow SM, Goloviznina NA, et al. Maternal high-fat diet and obesity compromise fetal hematopoiesis. Mol Metab 2015; 4(1): 25–38.

9. Farley D, Tejero ME, Comuzzie AG, et al. Feto-placental adaptations to maternal obesity in the baboon. Placenta 2009; 30(9): 752–60.

10. Gao X, Xu C, Asada N, Frenette PS. The hematopoietic stem cell niche: from embryo to adult. Development 2018; 145(2).

11. Bowie MB, Kent DG, Dykstra B, et al. Identification of a new intrinsically timed developmental checkpoint that reprograms key hematopoietic stem cell properties. Proc Natl Acad Sci U S A 2007; 104(14): 5878–82.

12. Morrison SJ, Hemmati HD, Wandycz AM, Weissman IL. The purification and characterization of fetal liver hematopoietic stem cells. Proc Natl Acad Sci U S A 1995; 92(22): 10302–6.

13. Coskun S, Chao H, Vasavada H, et al. Development of the fetal bone marrow niche and regulation of HSC quiescence and homing ability by emerging osteolineage cells. Cell Rep 2014; 9(2): 581–90.

14. Christensen JL, Wright DE, Wagers AJ, Weissman IL. Circulation and chemotaxis of fetal hematopoietic stem cells. PLoS Biol 2004; 2(3): E75.

15. Gomez Perdiguero E, Klapproth K, Schulz C, et al. Tissue-resident macrophages originate from yolk-sac-derived erythro-myeloid progenitors. Nature 2015; 518(7540): 547–51.

16. Hoeffel G, Chen J, Lavin Y, et al. C-Myb(+) erythro-myeloid progenitor-derived fetal monocytes give rise to adult tissue-resident macrophages. Immunity 2015; 42(4): 665–78.

17. Hoeffel G, Ginhoux F. Fetal monocytes and the origins of tissue-resident macrophages. Cell Immunol 2018; 330: 5–15.

18. Hoeffel G, Ginhoux F. Ontogeny of Tissue-Resident Macrophages. Front Immunol 2015; 6: 486.

19. McGrath KE, Frame JM, Palis J. Early hematopoiesis and macrophage development. Semin Immunol 2015; 27(6): 379–87.

20. Orkin SH, Zon LI. Hematopoiesis: an evolving paradigm for stem cell biology. Cell 2008; 132(4): 631–44.

21. Golub R, Cumano A. Embryonic hematopoiesis. Blood Cells Mol Dis 2013; 51(4): 226–31.

22. Duffield JS, Lupher M, Thannickal VJ, Wynn TA. Host responses in tissue repair and fibrosis. Annu Rev Pathol 2013; 8: 241–76.

23. Hornef MW, Fulde M. Ontogeny of intestinal epithelial innate immune responses. Front Immunol 2014; 5: 474.

24. Friedman JE. Developmental Programming of Obesity and Diabetes in Mouse, Monkey, and Man in 2018: Where Are We Headed? Diabetes 2018; 67(11): 2137–51.

25. Kim S, Kim N, Presson AP, et al. Dynamics of HSPC repopulation in nonhuman primates revealed by a decade-long clonal-tracking study. Cell Stem Cell 2014; 14(4): 473–85.

26. Sykes SM, Scadden DT. Modeling human hematopoietic stem cell biology in the mouse. Semin Hematol 2013; 50(2): 92–100.

27. Radtke S, Chan YY, Sippel TR, Kiem HP, Rongvaux A. MISTRG mice support engraftment and assessment of nonhuman primate hematopoietic stem and progenitor cells. Exp Hematol 2019; 70: 31–41 e1.

28. Wu C, Espinoza DA, Koelle SJ, et al. Geographic clonal tracking in macaques provides insights into HSPC migration and differentiation. J Exp Med 2018; 215(1): 217–32.

29. Messaoudi I, Estep R, Robinson B, Wong SW. Nonhuman primate models of human immunology. Antioxid Redox Signal 2011; 14(2): 261–73.

30. Pound LD, Kievit P, Grove KL. The nonhuman primate as a model for type 2 diabetes. Curr Opin Endocrinol Diabetes Obes 2014; 21(2): 89–94.

31. True CA, Takahashi DL, Burns SE, et al. Chronic combined hyperandrogenemia and western-style diet in young female rhesus macaques causes greater metabolic impairments compared to either treatment alone. Hum Reprod 2017; 32(9): 1880–91.

32. Bishop CV, Mishler EC, Takahashi DL, et al. Chronic hyperandrogenemia in the presence and absence of a western-style diet impairs ovarian and uterine structure/function in young adult rhesus monkeys. Hum Reprod 2018; 33(1): 128–39.

33. Bishop CV, Takahashi D, Mishler E, et al. Individual and combined effects of 5-year exposure to hyperandrogenemia and Western-style diet on metabolism and reproduction in female rhesus macaques. Hum Reprod 2021; 36(2): 444–54.

34. Varlamov O, Bishop CV, Handu M, et al. Combined androgen excess and Western-style diet accelerates adipose tissue dysfunction in young adult, female nonhuman primates. Hum Reprod 2017; 32(9): 1892–902.

35. Robino JJ, Pamir N, Rosario S, et al. Spatial and biochemical interactions between bone marrow adipose tissue and hematopoietic stem and progenitor cells in rhesus macaques. Bone 2020: 115248.

36. Thorn SR, Baquero KC, Newsom SA, et al. Early life exposure to maternal insulin resistance has persistent effects on hepatic NAFLD in juvenile nonhuman primates. Diabetes 2014; 63(8): 2702–13.

37. Varlamov O, Bucher M, Myatt L, Newman N, Grant KA. Daily Ethanol Drinking Followed by an Abstinence Period Impairs Bone Marrow Niche and Mitochondrial Function of Hematopoietic Stem/Progenitor Cells in Rhesus Macaques. Alcohol Clin Exp Res 2020.

38. Burwitz BJ, Reed JS, Hammond KB, et al. Technical advance: liposomal alendronate depletes monocytes and macrophages in the nonhuman primate model of human disease. J Leukoc Biol 2014; 96(3): 491–501.

39. Kowalczyk MS, Tirosh I, Heckl D, et al. Single-cell RNA-seq reveals changes in cell cycle and differentiation programs upon aging of hematopoietic stem cells. Genome Res 2015; 25(12): 1860–72.

40. Stuart T, Butler A, Hoffman P, et al. Comprehensive Integration of Single-Cell Data. Cell 2019; 177(7): 1888–902 e21.

41. Pellin D, Loperfido M, Baricordi C, et al. A comprehensive single cell transcriptional landscape of human hematopoietic progenitors. Nat Commun 2019; 10(1): 2395.

42. Carbone L, Davis BA, Fei SS, et al. Synergistic Effects of Hyperandrogenemia and Obesogenic Western-style Diet on Transcription and DNA Methylation in Visceral Adipose Tissue of Nonhuman Primates. Sci Rep 2019; 9(1): 19232.

43. Miller I, Min M, Yang C, et al. Ki67 is a Graded Rather than a Binary Marker of Proliferation versus Quiescence. Cell Rep 2018; 24(5): 1105–12 e5.

44. Kapellos TS, Bonaguro L, Gemund I, et al. Human Monocyte Subsets and Phenotypes in Major Chronic Inflammatory Diseases. Front Immunol 2019; 10: 2035.

45. Chang MY, Chan CK, Braun KR, et al. Monocyte-to-macrophage differentiation: synthesis and secretion of a complex extracellular matrix. J Biol Chem 2012; 287(17): 14122–35.

46. Asplund A, Stillemark-Billton P, Larsson E, et al. Hypoxic regulation of secreted proteoglycans in macrophages. Glycobiology 2010; 20(1): 33–40.

47. Asplund A, Friden V, Stillemark-Billton P, Camejo G, Bondjers G. Macrophages exposed to hypoxia secrete proteoglycans for which LDL has higher affinity. Atherosclerosis 2011; 215(1): 77–81.

48. Calvanese V, Nguyen AT, Bolan TJ, et al. MLLT3 governs human haematopoietic stem-cell self-renewal and engraftment. Nature 2019; 576(7786): 281–6.

49. Sugimura R, Jha DK, Han A, et al. Haematopoietic stem and progenitor cells from human pluripotent stem cells. Nature 2017; 545(7655): 432–8.

50. Lebert-Ghali CE, Fournier M, Kettyle L, Thompson A, Sauvageau G, Bijl JJ. Hoxa cluster genes determine the proliferative activity of adult mouse hematopoietic stem and progenitor cells. Blood 2016; 127(1): 87–90.

51. Jacob B, Osato M, Yamashita N, et al. Stem cell exhaustion due to Runx1 deficiency is prevented by Evi5 activation in leukemogenesis. Blood 2010; 115(8): 1610–20.

52. Ranzoni AM, Tangherloni A, Berest I, et al. Integrative Single-Cell RNA-Seq and ATAC-Seq Analysis of Human Developmental Hematopoiesis. Cell Stem Cell 2021; 28(3): 472–87 e7.

53. Comazzetto S, Shen B, Morrison SJ. Niches that regulate stem cells and hematopoiesis in adult bone marrow. Dev Cell 2021; 56(13): 1848–60.

54. Xu S, Lee KG, Huo J, Kurosaki T, Lam KP. Combined deficiencies in Bruton tyrosine kinase and phospholipase Cgamma2 arrest B-cell development at a pre-BCR+ stage. Blood 2007; 109(8): 3377–84.

55. Ng AP, Coughlan HD, Hediyeh-Zadeh S, et al. An Erg-driven transcriptional program controls B cell lymphopoiesis. Nat Commun 2020; 11(1): 3013.

56. Bergiers I, Andrews T, Vargel Bolukbasi O, et al. Single-cell transcriptomics reveals a new dynamical function of transcription factors during embryonic hematopoiesis. Elife 2018; 7.

57. Ding Y, Wang W, Ma D, et al. Smarca5-mediated epigenetic programming facilitates fetal HSPC development in vertebrates. Blood 2021; 137(2): 190–202.

58. Au WC, Moore PA, LaFleur DW, Tombal B, Pitha PM. Characterization of the interferon regulatory factor-7 and its potential role in the transcription activation of interferon A genes. J Biol Chem 1998; 273(44): 29210–7.

59. Singer K, DelProposto J, Morris DL, et al. Diet-induced obesity promotes myelopoiesis in hematopoietic stem cells. Mol Metab 2014; 3(6): 664–75.

60. Liu A, Chen M, Kumar R, et al. Bone marrow lympho-myeloid malfunction in obesity requires precursor cell-autonomous TLR4. Nat Commun 2018; 9(1): 708.

61. van den Berg SM, Seijkens TT, Kusters PJ, et al. Diet-induced obesity in mice diminishes hematopoietic stem and progenitor cells in the bone marrow. FASEB J 2016; 30(5): 1779–88.

62. Yan H, Baldridge MT, King KY. Hematopoiesis and the bacterial microbiome. Blood 2018; 132(6): 559–64.

63. Scheller EL, Doucette CR, Learman BS, et al. Region-specific variation in the properties of skeletal adipocytes reveals regulated and constitutive marrow adipose tissues. Nat Commun 2015; 6: 7808.

64. Rosen CJ, Ackert-Bicknell C, Rodriguez JP, Pino AM. Marrow fat and the bone microenvironment: developmental, functional, and pathological implications. Crit Rev Eukaryot Gene Expr 2009; 19(2): 109–24.

65. Ambrosi TH, Scialdone A, Graja A, et al. Adipocyte Accumulation in the Bone Marrow during Obesity and Aging Impairs Stem Cell-Based Hematopoietic and Bone Regeneration. Cell Stem Cell 2017; 20(6): 771–84 e6.

66. Li Z, Hardij J, Bagchi DP, Scheller EL, MacDougald OA. Development, regulation, metabolism and function of bone marrow adipose tissues. Bone 2018; 110: 134–40.

67. Adler BJ, Kaushansky K, Rubin CT. Obesity-driven disruption of haematopoiesis and the bone marrow niche. Nat Rev Endocrinol 2014; 10(12): 737–48.

68. Ernst LMR, E.D.; Carreon, C.K.; Huff, D.S.. Color Atlas of Human Fetal and Neonatal Histology: Springer; 2019.

69. Naveiras O, Nardi V, Wenzel PL, Hauschka PV, Fahey F, Daley GQ. Bone-marrow adipocytes as negative regulators of the haematopoietic microenvironment. Nature 2009; 460(7252): 259–63.

70. Butler JT, Abdelhamed S, Kurre P. Extracellular vesicles in the hematopoietic microenvironment. Haematologica 2018; 103(3): 382–94.

71. Huan J, Hornick NI, Goloviznina NA, et al. Coordinate regulation of residual bone marrow function by paracrine trafficking of AML exosomes. Leukemia 2015; 29(12): 2285–95.

72. Hornick NI, Doron B, Abdelhamed S, et al. AML suppresses hematopoiesis by releasing exosomes that contain microRNAs targeting c-MYB. Sci Signal 2016; 9(444): ra88.

73. Kalbermatter C, Fernandez Trigo N, Christensen S, Ganal-Vonarburg SC. Maternal Microbiota, Early Life Colonization and Breast Milk Drive Immune Development in the Newborn. Front Immunol 2021; 12: 683022.

74. Bauer SR, Kudo A, Melchers F. Structure and pre-B lymphocyte restricted expression of the VpreB in humans and conservation of its structure in other mammalian species. EMBO J 1988; 7(1): 111–6.

75. Shirasawa T, Ohnishi K, Hagiwara S, et al. A novel gene product associated with mu chains in immature B cells. EMBO J 1993; 12(5): 1827–34.

76. Goto T, Kennel SJ, Abe M, et al. A novel membrane antigen selectively expressed on terminally differentiated human B cells. Blood 1994; 84(6): 1922–30.

77. Gold MR, Matsuuchi L, Kelly RB, DeFranco AL. Tyrosine phosphorylation of components of the B-cell antigen receptors following receptor crosslinking. Proc Natl Acad Sci U S A 1991; 88(8): 3436–40.

78. Friedman JE, Dobrinskikh E, Alfonso-Garcia A, et al. Pyrroloquinoline quinone prevents developmental programming of microbial dysbiosis and macrophage polarization to attenuate liver fibrosis in offspring of obese mice. Hepatol Commun 2018; 2(3): 313–28.

79. Edlow AG, Glass RM, Smith CJ, Tran PK, James K, Bilbo S. Placental Macrophages: A Window Into Fetal Microglial Function in Maternal Obesity. Int J Dev Neurosci 2019; 77: 60–8.

80. Russo L, Muir L, Geletka L, et al. Cholesterol 25-hydroxylase (CH25H) as a promoter of adipose tissue inflammation in obesity and diabetes. Mol Metab 2020; 39: 100983.

81. Reilly SM, Saltiel AR. Adapting to obesity with adipose tissue inflammation. Nat Rev Endocrinol 2017; 13(11): 633–43.

82. Singer K, Maley N, Mergian T, et al. Differences in Hematopoietic Stem Cells Contribute to Sexually Dimorphic Inflammatory Responses to High Fat Diet-induced Obesity. J Biol Chem 2015; 290(21): 13250–62.

83. McNelis JC, Olefsky JM. Macrophages, immunity, and metabolic disease. Immunity 2014; 41(1): 36–48.

84. Ho YH, Mendez-Ferrer S. Microenvironmental contributions to hematopoietic stem cell aging. Haematologica 2020; 105(1): 38–46.

85. Morrison SJ, Wandycz AM, Akashi K, Globerson A, Weissman IL. The aging of hematopoietic stem cells. Nat Med 1996; 2(9): 1011–6.

86. Netea MG, Dominguez-Andres J, Barreiro LB, et al. Defining trained immunity and its role in health and disease. Nat Rev Immunol 2020; 20(6): 375–88.

87. Jonscher KR, Abrams J, Friedman JE. Maternal Diet Alters Trained Immunity in the Pathogenesis of Pediatric NAFLD. J Cell Immunol 2020; 2(6): 315–25.

