## Supplementary material for "Maternal Western-Style Diet Impairs Bone Marrow Development and Drives a Hyperinflammatory Phenotype in Hematopoietic Stem and Progenitor Cells in Fetal Rhesus Macaques": Supp Figures

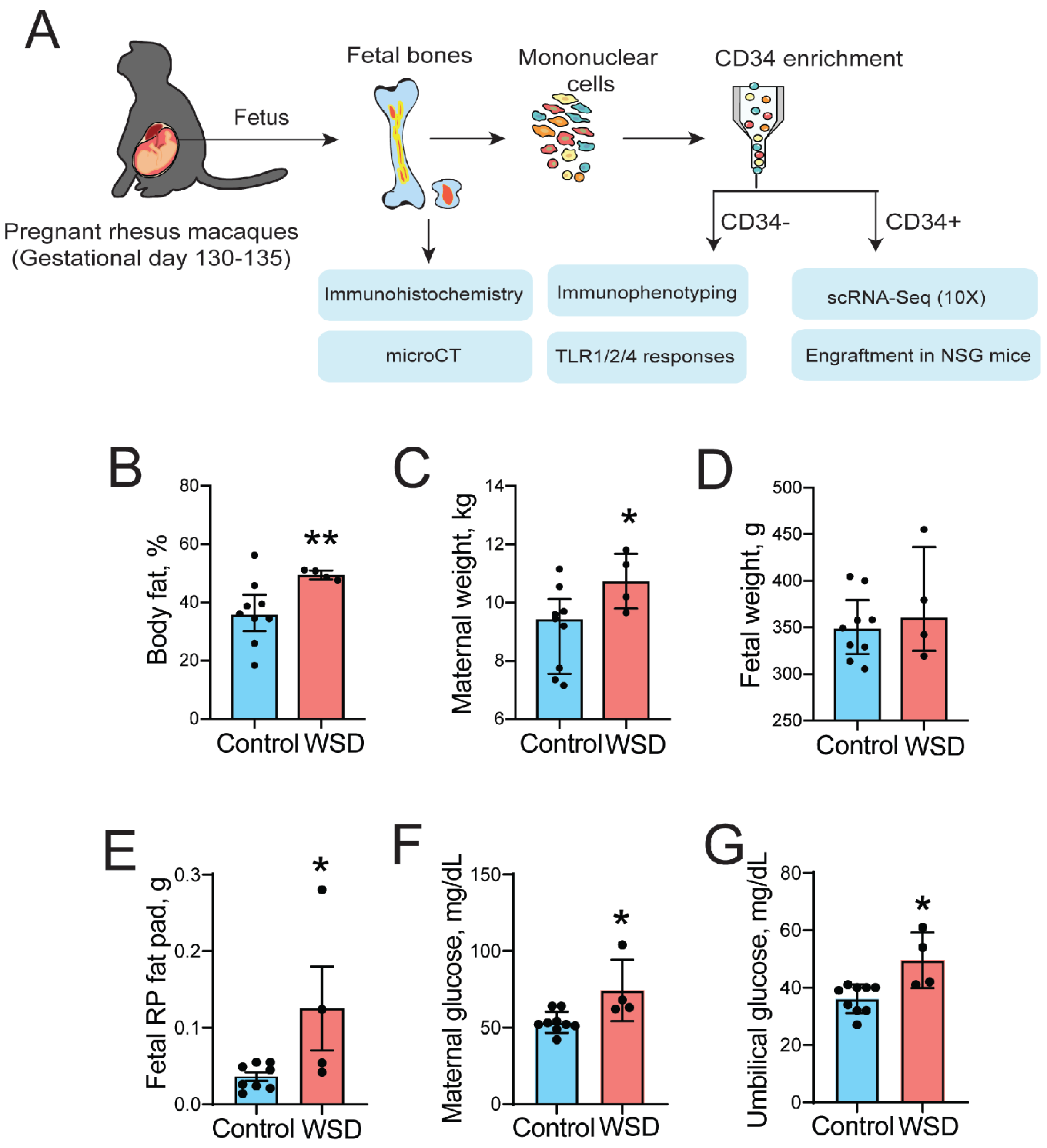


**Supplementary Figure S1. Experimental design.** (A) Experimental approach; long bones (tibias and femurs) and flat bones (sterna) from gestational day 130-135 fetuses were isolated from dams on a control diet and on a WSD. Femurs and sterna were used for immunohistochemistry and histology. Femurs were scanned using micro-computed tomography (microCT). Tibias were crushed and CD34+ HSPCs were immuno-isolated from the mononuclear fraction using magnetic bead. CD34+ cells were enrichment and profiled using scRNA-Seq and engrafted in nonlethally irradiated NSG mice. Tibial mononuclear cells from the CD34-negative fraction was phenotyped using flow cytometry and functional responses to TLR ligand stimulation were tested. (B) Maternal body fat before pregnancy, (C) maternal weights at C-section, (D) fetal weights, (E) Fetal retroperitoneal (RP) fat pad weight at C-section, (F) maternal and (G) umbilical artery glucose levels at C-section. Bars are means ± SEM; t-test *$p<0.05$, **$p<0.01$.

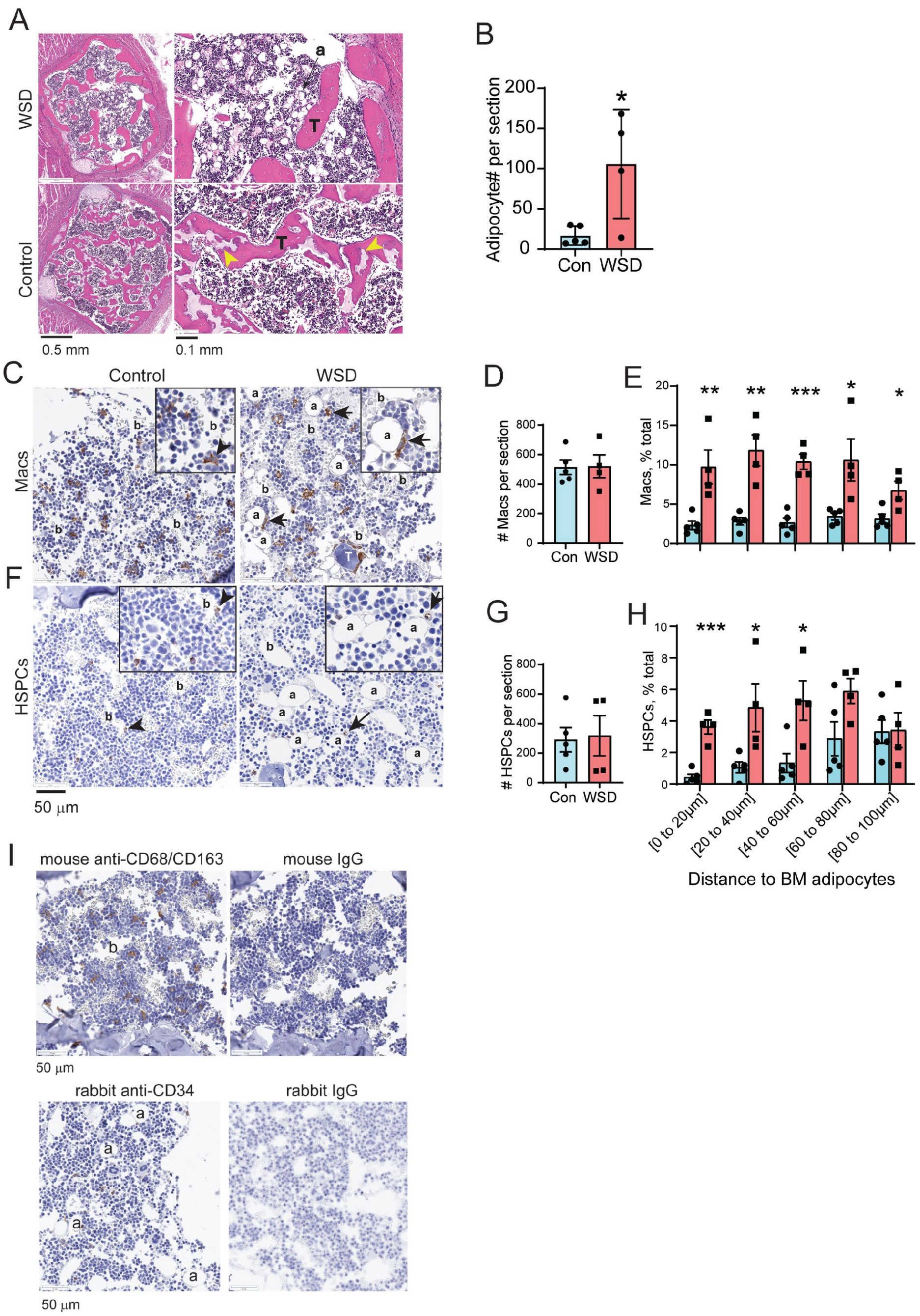


**Supplementary Figure S2. Immunohistochemical analysis of fetal sterna.** (A) Representative images of H&E stained sternum from the control and WSD fetuses; a, adipocytes; T, trabeculae; yellow arrowheads, cartilaginous areas of partially ossified trabeculae. (B) Adipocyte content in fetal sterna; bars are means ± SEM;*p<0.05; each data point represents one fetus. (C and F) Representative images of fetal sterna from the control and WSD fetuses stained with anti-CD68/CD163 (macrophages, Macs) or anti-CD34 (HSPCs) antibodies. Macs and HSPCs located near BM adipocytes are marked with arrows; b, identifiable blood vessel filled with red blood cells. Insets, enlarged images showing close contacts between adipocytes and Macs/HSPCs. (D) Mac and (G) HSPC content in fetal sterna; the total number of cells per sternal section is shown. The % of total Macs (E) and HSPCs (H) within indicated distances to FBM adipocytes. Bars are means ± SEM. All two-group comparisons were tested for statistical differences using unpaired t-tests; *p<0.05; **p<0.01; ***p<0.001. (I) Evaluation of antibodies to CD68/CD68 and CD34 using fetal sterna. Negative controls included isotype-specific antibodies.

A

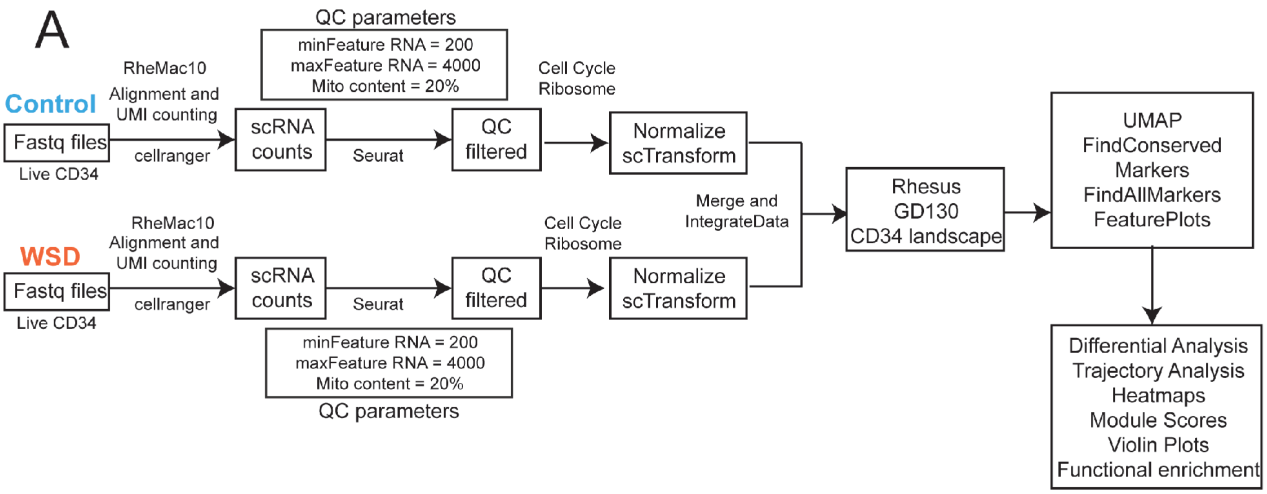


B

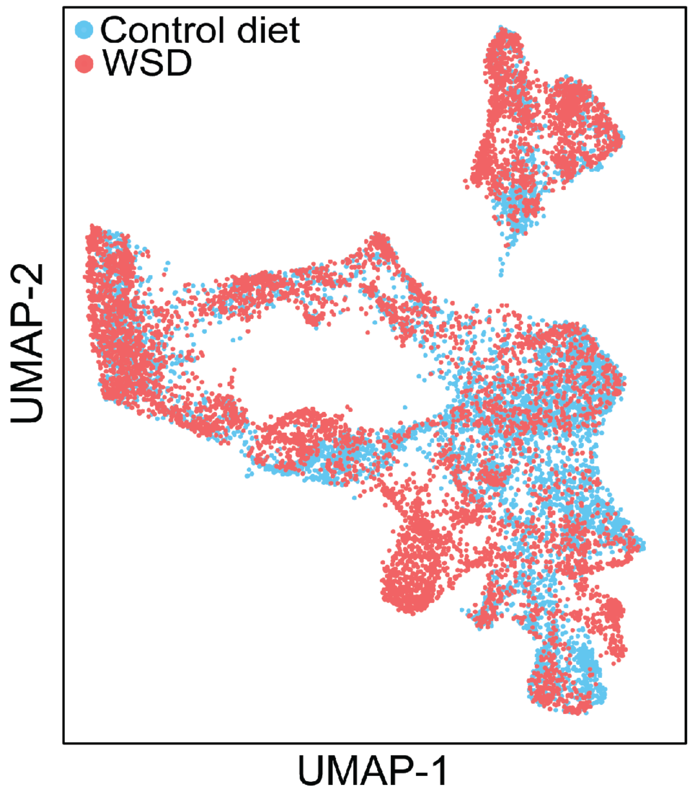


**Supplementary Figure S3: Single cell analysis of CD34+ progenitors in the FBM.** (A) Analysis strategy for scRNA-Seq data. (B) UMAP representation of fetal bone marrow CD34+ cells highlighting differences with maternal diet.

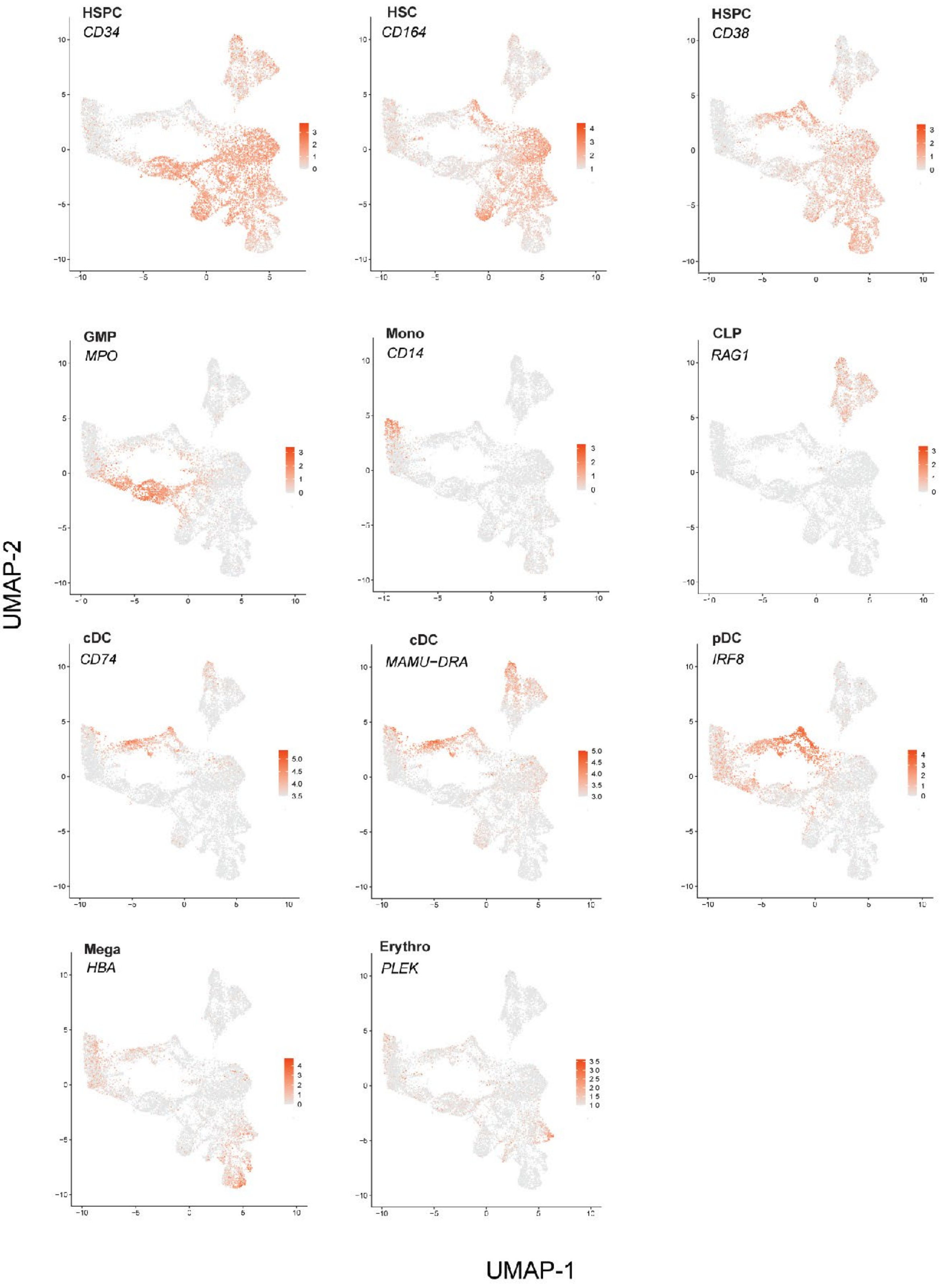


**Supplementary Figure S4. Lineage-specific gene expression in FBM HSPCs.** Feature plots highlighting markers expressed in hematopoietic progenitor populations in FBM HSPCs. Progenitor types are indicated.

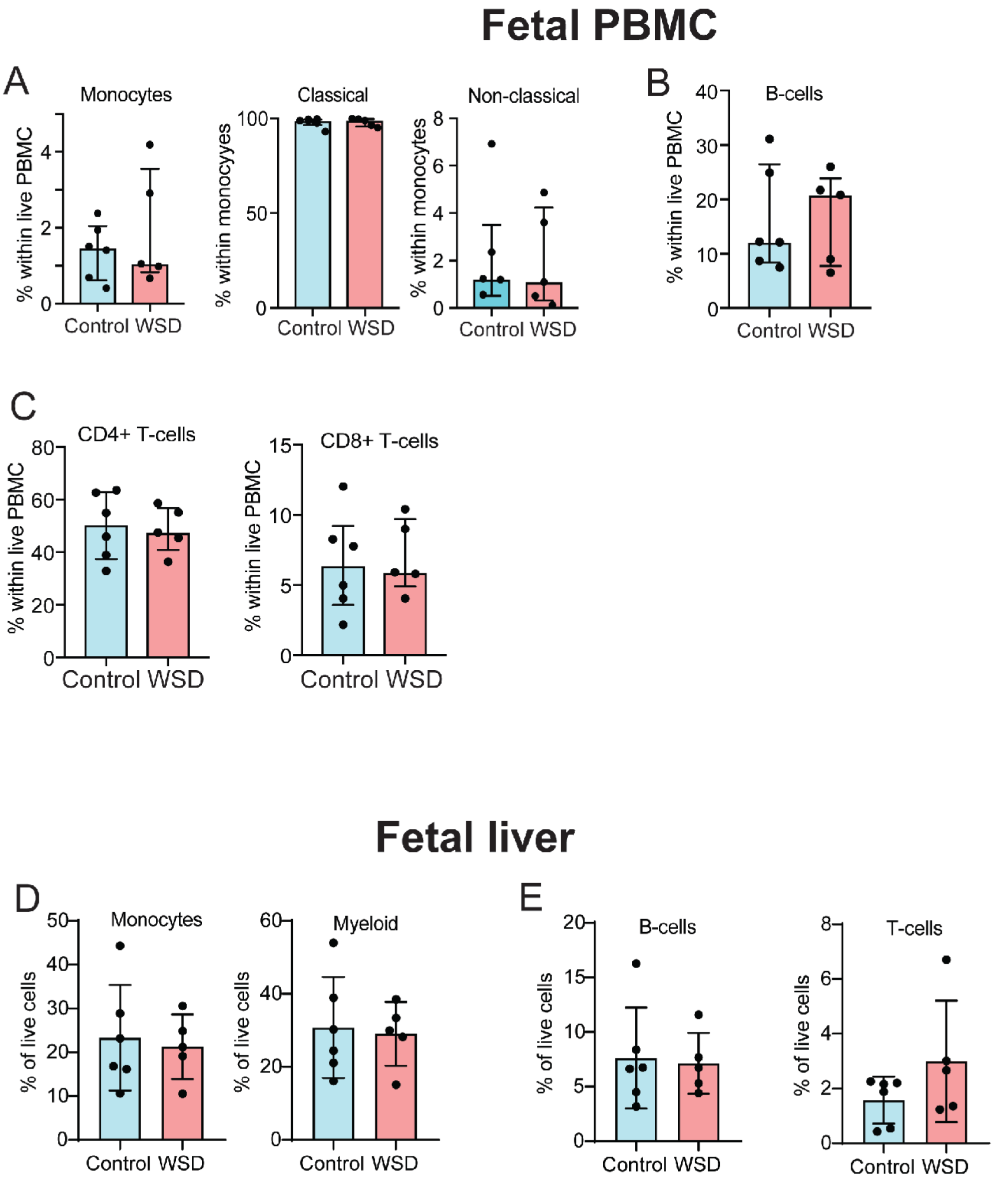


**Supplementary Figure S5. Effect of maternal diet on peripheral blood and fetal liver immune subsets.** Relative frequencies of (A) CD14+ monocytes and its subsets (classical CD14+CD16-); non-classical CD14+CD16+), (B) CD20+ B-cells, and (C) T-cell subsets in fetal peripheral blood. Relative frequencies of (D) CD11b+CD14+ monocytes and CD11b+ myeloid cells, (E) CD20+ B-cells and CD3+ T-cells in the fetal liver.

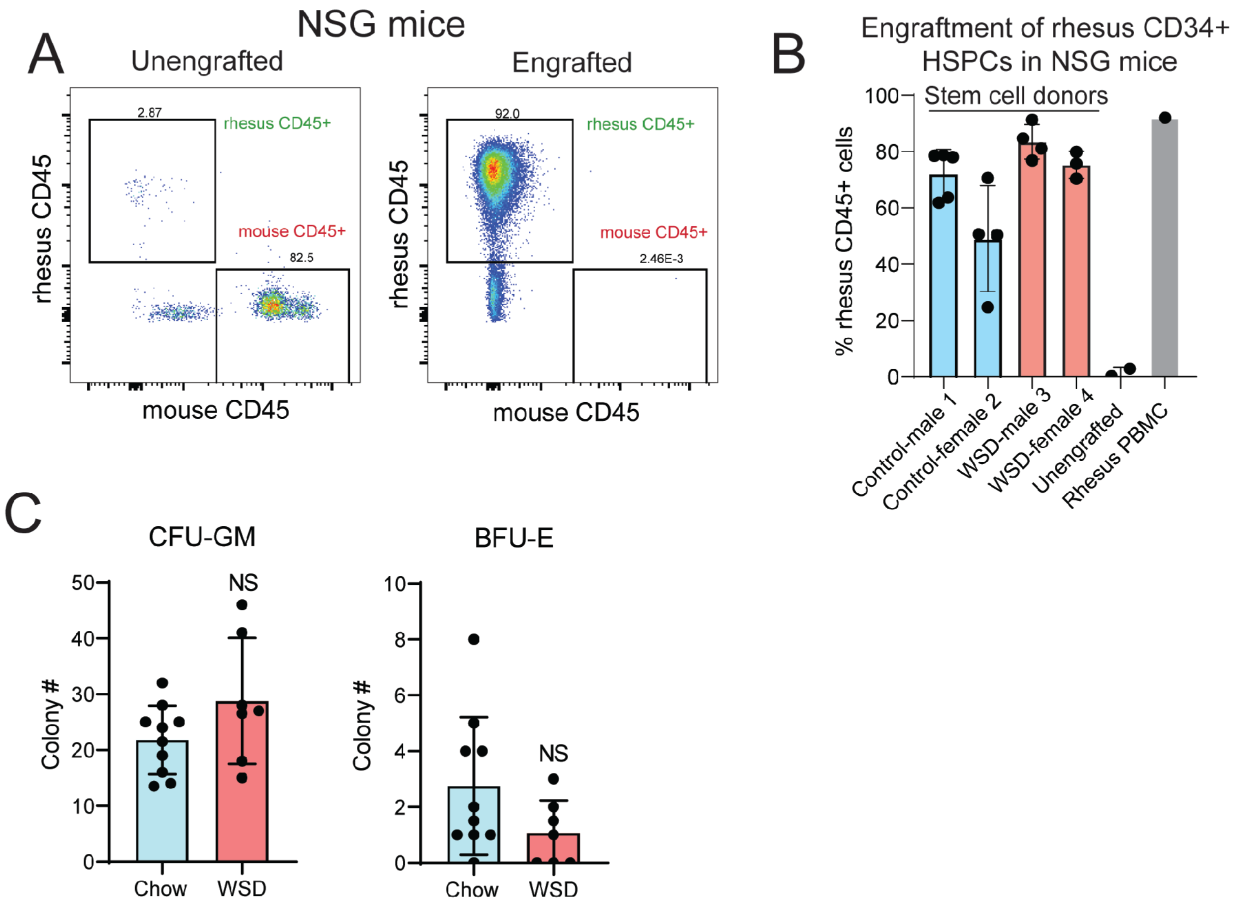


**Supplementary Figure S6. The engraftment analysis of FBM HSPCs in NSG mice.** (A) Example of flow cytometry analysis of peripheral blood from an NSG mouse engrafted with fetal rhesus CD34+ cells in comparison with unengrafted controls. (B) Relative frequencies of rhesus CD45+ cells in the peripheral blood of NSG mice engrafted with rhesus CD34+ cells in comparison with unengrafted controls. Rhesus PBMC, positive staining control. (C) In vitro colony-forming (CFU) assay of rhesus CD34+ cells isolated from the tibial BM of NSG mice. Rhesus HSPCs were isolated using NHP-specific antibodies to CD34 and 250 cells were plated on MethoCult™ (H4435 Enriched) supplied with 100 ng/ml SCF, TPO, and Flt3L. CFU-GM and BFU-E colonies were scored after 10 days in culture. Bars are means ± SEM. Each biological data point represents one NGG mouse (the mean of three replicate experiments). NS, not significant.
